# The minimum land area requiring conservation attention to safeguard biodiversity

**DOI:** 10.1101/839977

**Authors:** James R. Allan, Hugh P. Possingham, Scott C. Atkinson, Anthony Waldron, Moreno Di Marco, Vanessa M. Adams, Stuart H. M. Butchart, W. Daniel Kissling, Thomas Worsdell, Gwili Gibbon, Kundan Kumar, Piyush Mehta, Martine Maron, Brooke A. Williams, Kendall R. Jones, Brendan A. Wintle, April E. Reside, James E.M. Watson

**Affiliations:** Institute for Biodiversity and Ecosystem Dynamics (IBED), University of Amsterdam, P.O. Box 94240, 1090 GE, Amsterdam, The Netherlands; Centre for Biodiversity and Conservation Science, The University of Queensland, St Lucia, QLD 4072, Australia; The Nature Conservancy, VA 22203-1606, USA; United Nations Development Programme (UNDP), New York, New York, USA; Cambridge Conservation Initiative, David Attenborough Building, Department of Zoology, Cambridge University, Cambridge CB2 3QZ, UK; Department of Biology and Biotechnologies, Sapienza University of Rome, viale dell’Università 32, I-00185 Rome, Italy; School of Earth and Environmental Sciences, The University of Queensland, St Lucia QLD 4072, Australia; School of Technology, Environments & Design, University of Tasmania, Hobart, TAS 7001, Australia; BirdLife International, David Attenborough Building, Pembroke Street, Cambridge CB2 3QZ, UK; Department of Zoology, University of Cambridge, Downing Street, Cambridge CB2 3EJ, UK; Rights and Resources Initiative, Washington, D. C., USA; Durrell Institute of Conservation and Ecology, School of Anthropology and Conservation, University of Kent, Canterbury, CT2 7NR, UK; University of Delaware, Newark, DE 19716, USA; Wildlife Conservation Society, Global Conservation Program, 2300 Southern Boulevard, Bronx, NY 10460-1068, USA; School of BioSciences, University of Melbourne, Vic., Australia

**Keywords:** Protected Areas, Conservation Planning, Restoration, Global Conservation Priorities, Strategic Plan for Biodiversity, Convention on Biological Diversity, Conservation Priorities, Key Biodiversity Areas, Wilderness, Ecologically Intact

## Abstract

More ambitious conservation efforts are needed to stop the global biodiversity crisis. Here, we estimate the minimum land area to secure important sites for terrestrial fauna, ecologically intact areas, and the optimal locations for representation of species ranges and ecoregions. We discover that at least 64 million km^2^ (44% of terrestrial area) requires conservation attention. Over 1.8 billion people live on these lands so responses that promote agency, self-determination, equity, and sustainable management for safeguarding biodiversity are essential. Spatially explicit land-use scenarios suggest that 1.3 million km^2^ of land requiring conservation could be lost to intensive human land-uses by 2030, which requires immediate attention. However, there is a seven-fold difference between the amount of habitat converted under optimistic and pessimistic scenarios, highlighting an opportunity to avert this crisis. Appropriate targets in the post-2020 Global Biodiversity Framework to ensure conservation of the identified land would contribute substantially to safeguarding biodiversity.

---

Securing places with high conservation value is crucial for safeguarding biodiversity^1^and is central to the Convention on Biological Diversity (CBD)’s 2050 vision of sustaining a healthy planet and delivering benefits for all people^2^. CBD Aichi Target 11 aimed to conserve at least 17% of land area by 2020^3^, but this is widely seen as inadequate for halting biodiversity declines and averting the crisis^4^. Post-2020 target discussions are now well underway^5^, and there is a broad consensus that the amount of land and sea managed for biodiversity conservation must increase^6^. Recent calls are for targets to conserve anywhere from 26 to 60% of land and ocean area by 2030 through site-scale responses such as protected areas and ‘other effective area-based conservation measures’ (OECMs)^7-12^. There is also increasing recognition that site-scale responses must be supplemented by broader landscape-scale actions aimed at addressing habitat degradation and loss^13^. While global conservation targets are set by political intergovernmental negotiation, scientific input is necessary to identify the location and amount of land requiring conservation attention, and to inform potential strategies.

Several scientific approaches exist that help provide evidence to inform global conservation efforts, but when used in isolation, they can provide conflicting advice. In particular, there are efficiency-based planning approaches that focus on maximising the number of species or ecosystems captured within a complementary set of conservation areas, weighting species and ecosystems by their endemicity, extinction risk, or other criteria^14,15^. There are also threshold-based approaches such as the Key Biodiversity Area (KBA) initiative^16^, which identifies sites of significance for the global persistence of biodiversity using criteria relating to the occurrence of threatened or geographically restricted species or ecosystems, intact ecological communities, or important biological processes (e.g. breeding aggregations)^16^. There are also proactive approaches that aim to conserve the most ecologically intact places before they are degraded^17^. These intact areas are increasingly recognised as essential for sustaining long-term ecological and evolutionary processes^18^, and long-term species persistence^19^, especially under climate change^20^. Examples include boreal forests which support many wide-ranging species^21,22^, and the Amazon rainforest which needs to be maintained in its entirety, not just for its most species-rich areas but also to sustain continent-scale hydrological patterns that underpin its ecosystems^23^.

Although these approaches are complementary and provide essential evidence to set and meet biodiversity conservation targets, the adoption of any one of them as a unique guide for decision-making is likely to omit potentially critical elements of the CBD vision^24^. For example, a species-based focus on identifying areas in a way that most efficiently captures the most species would fail to recognise the critical need to maintain large intact ecosystems globally for biodiversity persistence^19^. Equally, a focus on proactively conserving ecologically intact ecosystems would fail to achieve adequate conservation of some threatened species or ecosystems^25^. Put simply, all approaches will lead to partly overlapping but often distinct science-based suggestions for area-based conservation^26^. We suggest that combining these approaches into a unified global framework that seeks to comprehensively conserve species, ecosystems, and the remaining intact ecosystems offers a better scientific basis for achieving the CBD vision.

Here, we identify the minimum land area requiring conservation attention globally. We start from the basis of existing protected areas^27^, KBAs^28^, and ecologically intact areas^29^, and then efficiently add a fraction of the ranges of 35,561 species of mammals, birds, amphibians, reptiles, freshwater crabs, shrimp, and crayfish scaled to the sizes of their ranges^14,15,30^, while also capturing samples (17% of area, following CBD Aichi Target 11) of all terrestrial ecoregions^31^. We used these taxonomic groups because they are those most comprehensively assessed and mapped by the International Union for the Conservation of Nature (IUCN), noting that the inclusion of plants and other groups would likely increase the area identified above our minimum.

We do not suggest the land we map should be designated as protected areas that preclude other land management strategies. Rather, we argue that it should be managed through a range of strategies for species and ecosystem conservation. We define the term ‘conservation attention’ to capture this broad range of strategies which lead to positive biodiversity outcomes. For example, extensive areas that are remote and unlikely to be converted for human uses in the near-term could be safeguarded through effective sustainable land-use policies, while other areas could be conserved through self-determined local governance regimes led by Indigenous Peoples and Local Communities. We believe the appropriate governance and management regimes for any area depends in part on the likelihood of its habitat being converted or degraded by intensive human uses^32-34^ as well as the land tenure regimes and other socio-political factors and as such, the response for conserving the areas we identify will be context specific.

To highlight places that need the most immediate attention, we further calculate which parts of the land needing conservation are most likely to suffer habitat conversion in the near future. We do this by using harmonised projections of future land-use change by 2030 and 2050^35^. To determine best-to worst-case scenarios, we evaluated projections under three different shared socioeconomic pathways (SSPs)^36^ linked to representative concentration pathways (RCPs)^37^: an optimistic scenario where the world gradually moves towards a more sustainable future, SSP1 (RCP2.6; IMAGE model), a middle-of-the-road scenario without any extreme changes towards or away from sustainability (SSP2; MESSAGE-GLOBIOM model), and a pessimistic scenario where regional rivalries dominate international relations and land-use change is poorly regulated, SSP3 (RCP7.0; AIM model). Given uncertainty in which pathway humanity is following we also created an “ensemble” land-use projection where we calculated the average loss across all three SSPs.

We also estimate and map the number of people living on the land area requiring conservation attention, including within current protected areas, using the LandScan 2018 global distribution^38^. We performed this calculation in view of the potential impact of conservation on people living in such areas given the history of human rights abuses^39^, displacement^40^, and militarised forms of violence^41^ associated with some actions done in the name of conservation^42^. These rights-abuses are linked to a pervasive lack of tenure-rights recognition and culturally appropriate rights frameworks for conservation^43-45^. Communities already effectively conserve large tracts of land, and supporting their actions will thus be a key strategy to continue safeguarding biodiversity^46^.

## The minimum land area requiring conservation attention

We estimate that, in total, the minimum land area requiring conservation attention is 64.7 million km^2^ (44% of Earth’s terrestrial area; Figure 1). This consists of 35.1 million km^2^ of ecologically intact areas, 20.5 million km^2^ of existing protected areas, 11.6 million km^2^ of KBAs, and 12.4 million km^2^ (8.4% of terrestrial area) of additional land (i.e. outside protected areas, KBAs and ecologically intact areas) needed to promote species persistence based on conserving minimum proportions of their ranges (Figure 2). Moreover, protected areas, KBAs and ecologically intact areas only have a three way overlap on 1.8 million km^2^, and consensus area (overlap) only captures 5% of ecologically intact areas, 9% of protected area extent, and 16% of KBA extent, emphasising the importance of considering the various approaches in a unified framework as we do here.

**Figure 1.**
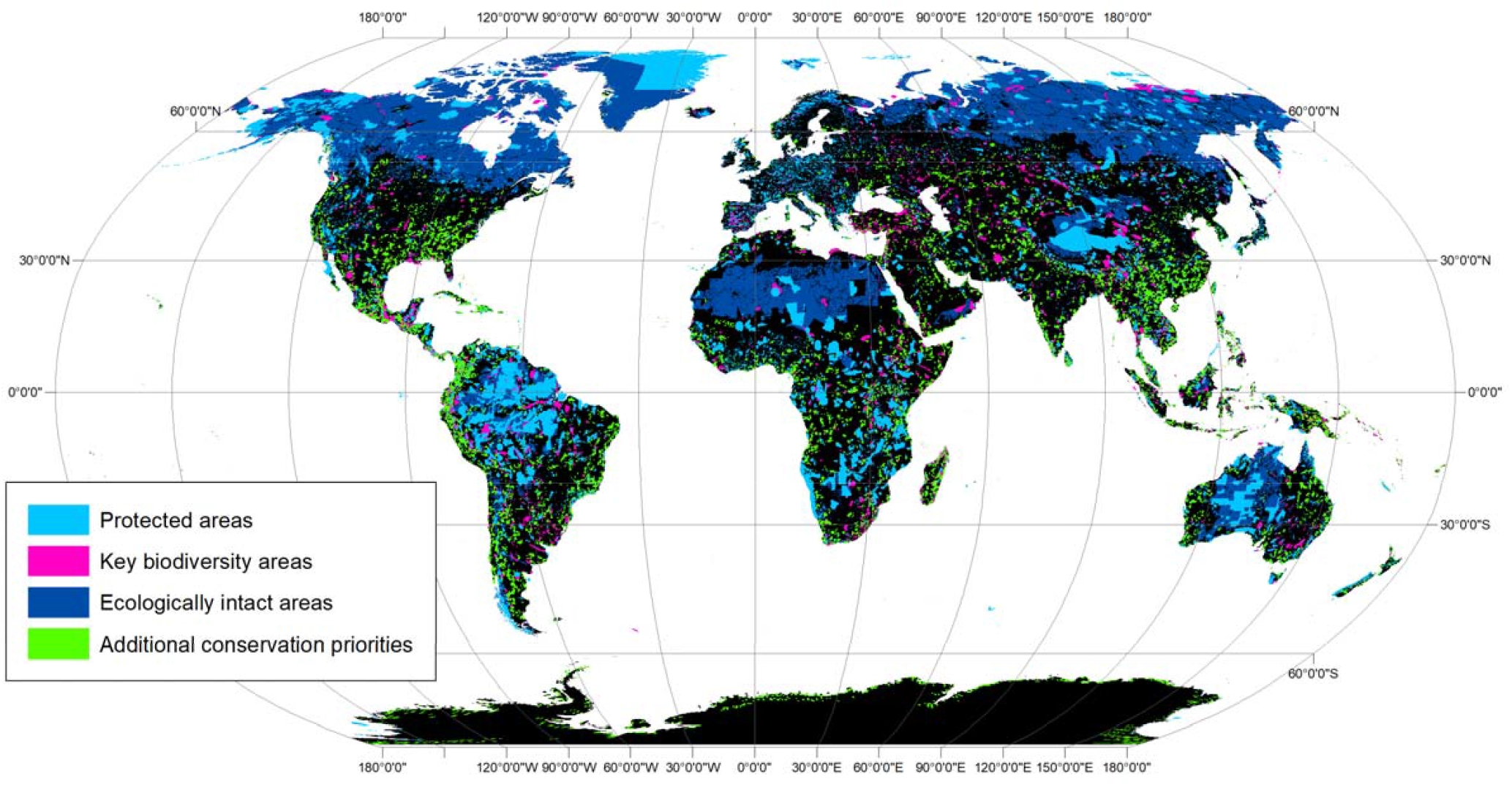
The minimum land area for conserving terrestrial biodiversity. The components include protected areas (light blue), Key Biodiversity Areas (purple) and ecologically intact areas (dark blue). Where they overlap, protected areas are shown above Key Biodiversity Areas, which are shown above ecologically intact areas. New conservation priorities are in green. The Venn diagram shows the proportional overlap between features.

**Figure 2.**
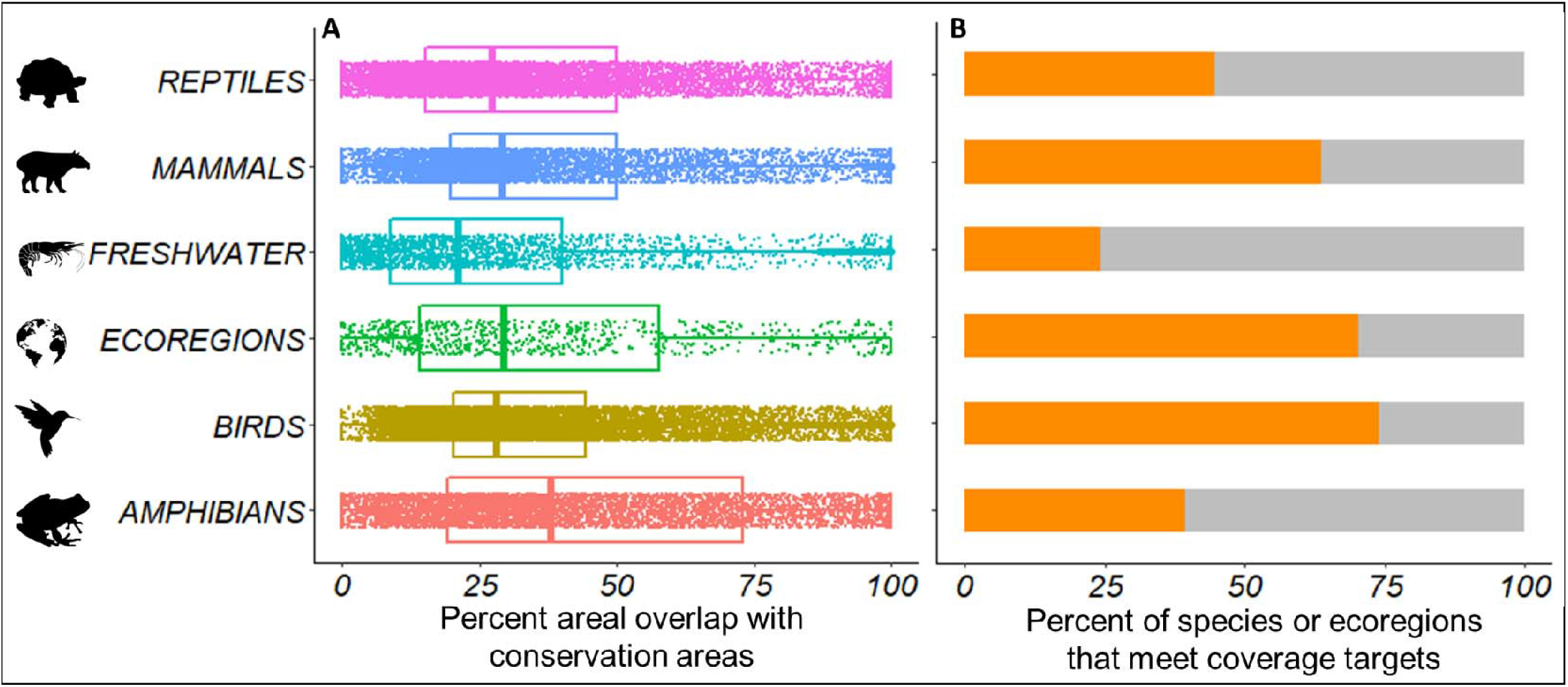
Gap analyses of species and ecoregion coverage within areas of conservation importance. A) The percentage of the distribution of each species (in different taxonomic groups; freshwater includes crabs, shrimp and crayfish) and ecoregion area that overlaps with areas of conservation importance (protected areas, Key Biodiversity Areas, and ecologically intact areas). Boxplots show the median and 25^th^ and 75^th^ percentiles for each taxonomic group. B) The percentage of species and ecoregions with an adequate proportion of their distribution overlapping existing conservation areas to meet specific coverage targets for species (10–100% depending on range size) or ecoregions (17%) (orange).

There is considerable geographic variation in the amount of land requiring conservation. We find that at least 64% of land in North America needs to be conserved, primarily due to the ecologically intact areas of Canada and the USA and extensive additional land areas in Central America. In contrast, at least 33.1% of Europe’s land area requires conservation. The proportion of land requiring conservation also varies considerably among nations (Figure 3), with notably high values in Canada (84%) largely due to its extensive ecologically intact areas, Costa Rica (86%), Suriname (84%), and Ecuador (81%), due to their high numbers of endemic species and, in Ecuador’s case, the inclusion of a large overlap with the remaining Amazon forest (Extended Data Table 1). We also find that a larger percentage of land in developed economies (55% in total) requires effective conservation compared to emerging economies (48%) or developing economies (30%) (Extended Data Table 2).

**Figure 3.**
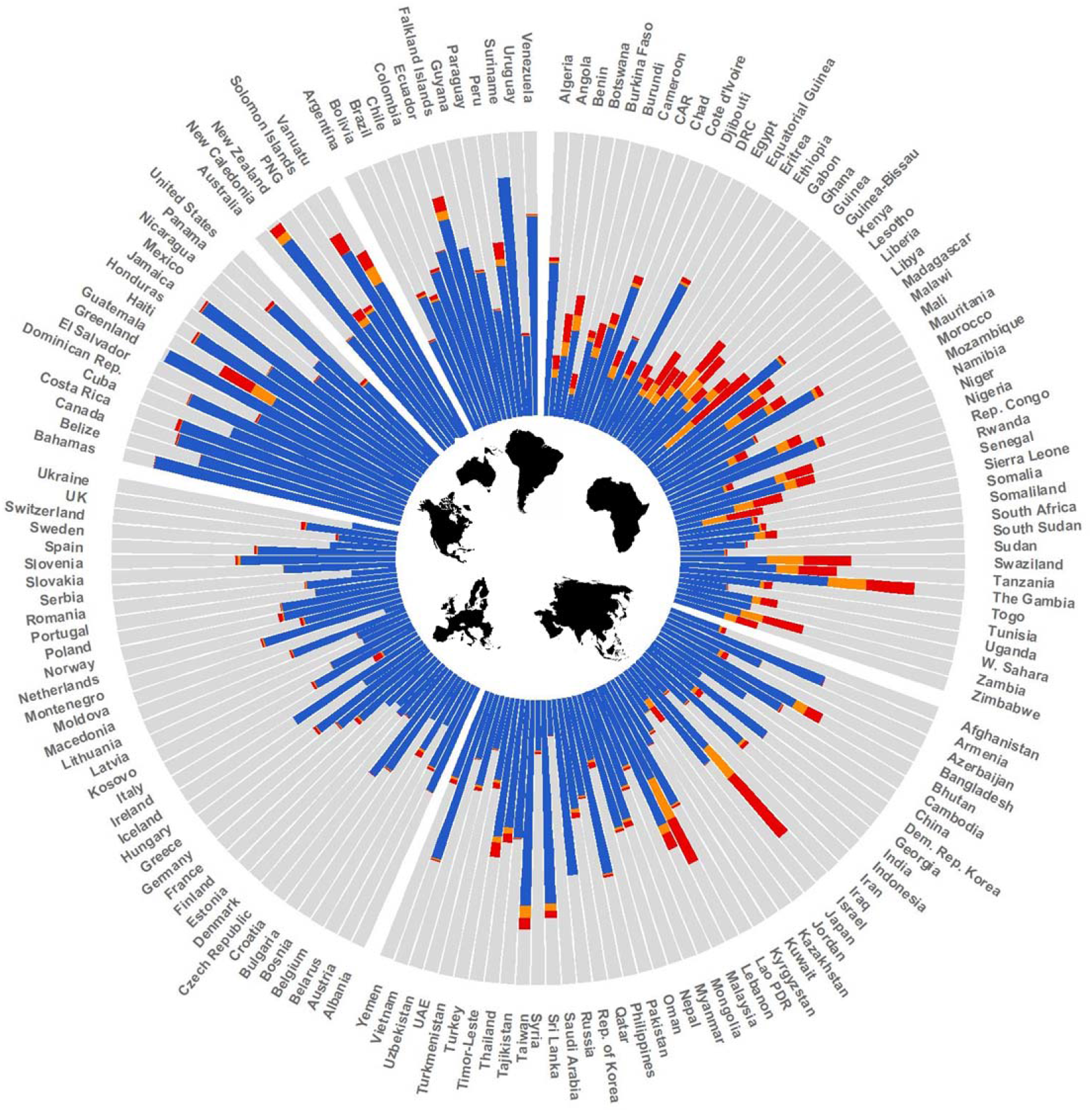
National level land area for conservation and projected habitat loss. Estimated proportion of each country requiring effective conservation attention that is projected to suffer habitat conversion by 2030 (orange) and 2050 (red) or that are projected not to be converted (blue) according to Shared Socioeconomic Pathway 3 (SSP3; a worst-case scenario). Grey areas are outside the land identified for conservation. We excluded 85 countries with a land area < 10,000 km^2^ from the figure.

## Future risk of land conversion in areas requiring conservation attention

We found that 44.9 million km^2^ (70.1%) of the land area requiring conservation attention is currently intact, implying a significant restoration requirement. Our results further suggest that under the pessimistic scenario SSP3, 1.3 million km^2^ (2.8%) of the total intact land area requiring conservation will undergo habitat conversion to intensive human land-uses by 2030, increasing to 2.2 million km^2^ (4.9%) by 2050. Projected habitat conversion varies across continents and countries (Figure 4). Africa is projected to have the highest proportion of intact conservation land converted by 2030 (>800,506 km^2^, 9% of Africa’s intact habitat), increasing to 1.4 million km^2^ (15.9%) by 2050 (Extended Data Tables 3-4). The lowest risk of conversion is in Oceania and North America. Substantially larger proportions of intact land requiring conservation in developing economies are projected to have their habitat converted by 2030 (7.1%), compared with emerging economies (1.7%) or developed economies (1.1%). By 2050, developing economies are projected to have 12.7% of their intact habitat requiring conservation converted under SSP3 (Extended Data Table 5), notably a lot of this loss is driven by demand in developed economies^47^. KBA’s are projected to have the largest proportion of habitat converted compared with protected areas and ecologically intact areas (Extended Data Table 6).

**Figure 4.**
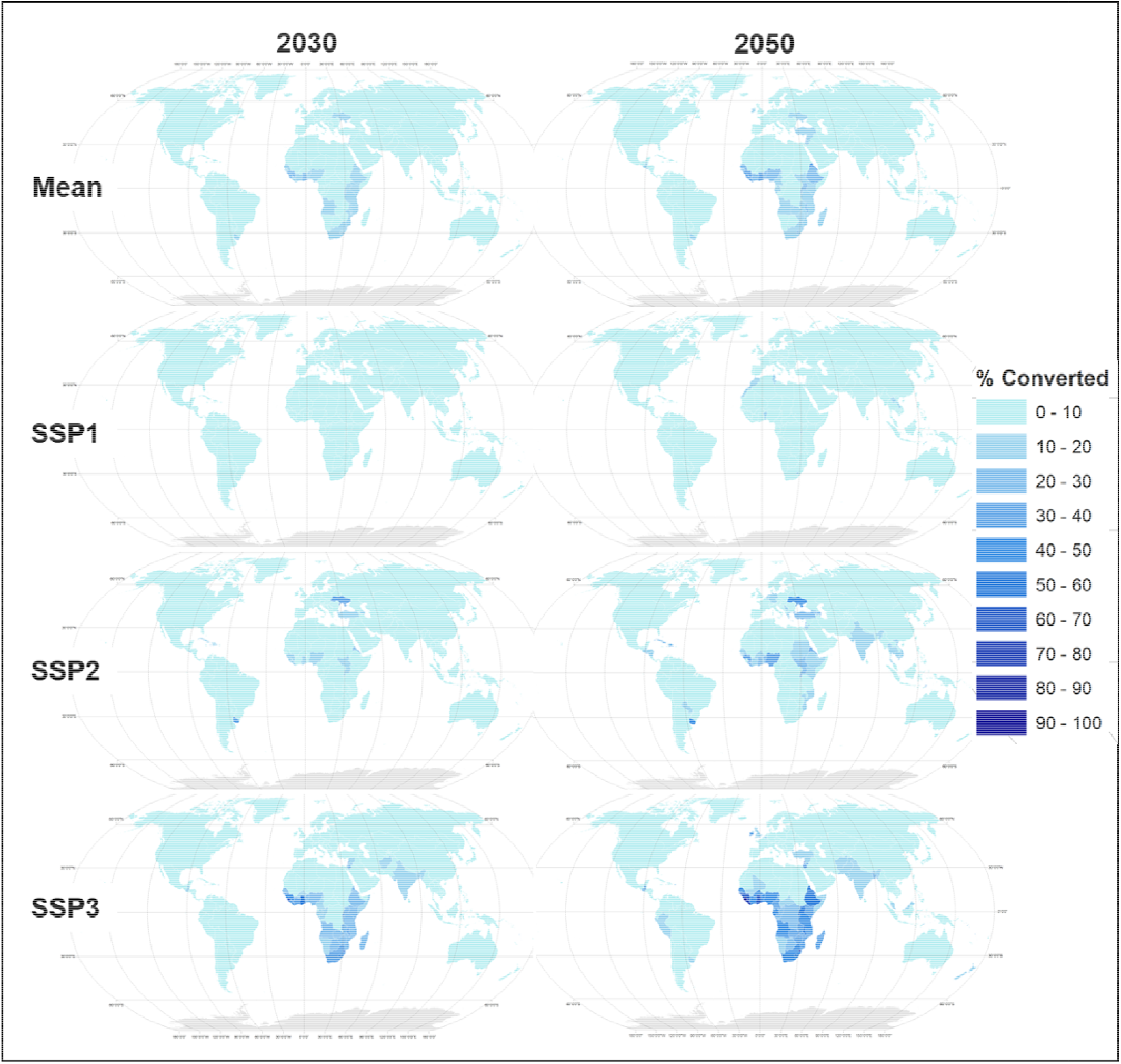
Future habitat conversion on land requiring conservation attention. The proportion of natural habitat on land requiring conservation that is projected to be converted to human uses by 2030 and 2050 based on Shared Socioeconomic Pathway 1 (SSP1; an optimistic scenario), Shared Socioeconomic Pathway 2 (SSP2; a middle-of-the-road scenario), Shared Socioeconomic Pathway 3 (SSP3; a pessimistic scenario), and the mean loss across the three scenarios (Mean). The data on future land use does not extend to Antarctica.

Based on SSP1, representing a world acting sustainably, we estimate that 136,380 km^2^ (0.3%) of the intact land requiring effective conservation may suffer natural habitat conversion by 2030, increasing to 320,558 km^2^ (0.7%) by 2050. Based on SSP2, representing a middle-of-the-road scenario, the values become 841,438 km^2^ (1.9%) by 2030 and 1.5 million km^2^ (3.3%) by 2050. This highlights how our results are sensitive to future societal development pathways, but even under the most optimistic scenario (SSP1), large extents of important conservation land are at risk of having natural habitat converted to more intensive human land-uses. However, the seven-fold difference between the amount of habitat converted under SSP1 vs. SSP3 shows there is a large window of opportunity for humanity to avert the biodiversity crisis.

There is inherent uncertainty in future land-use projections and on which SSP society is tracking most closely. To minimise the effect of this uncertainty, we also calculated the average intact habitat loss across the three SSP scenarios. In this ‘ensemble’ scenario, we expect 740,599 km^2^ (1.7%) of intact habitat in land requiring conservation to be converted by 2030, increasing to 1.3 million km^2^ by 2050 (2.9%).

## Human population in areas requiring conservation

We found that 1.87 billion people live in the land area requiring conservation attention, which is approximately one-quarter of Earth’s human population (24%) (Extended Data Figure 1). Africa, Asia and Central America have particularly large proportions of their human populations living on important conservation land (Extended Data Figure 2). The majority of people living in the area requiring conservation are in emerging and developing economies, which also have much higher proportions of their populations (often above 20%) living in areas requiring conservation compared to developed economies (Figure 5). This raises social justice questions regarding scaling up conservation strategies to meet biodiversity goals.

**Figure 5.**
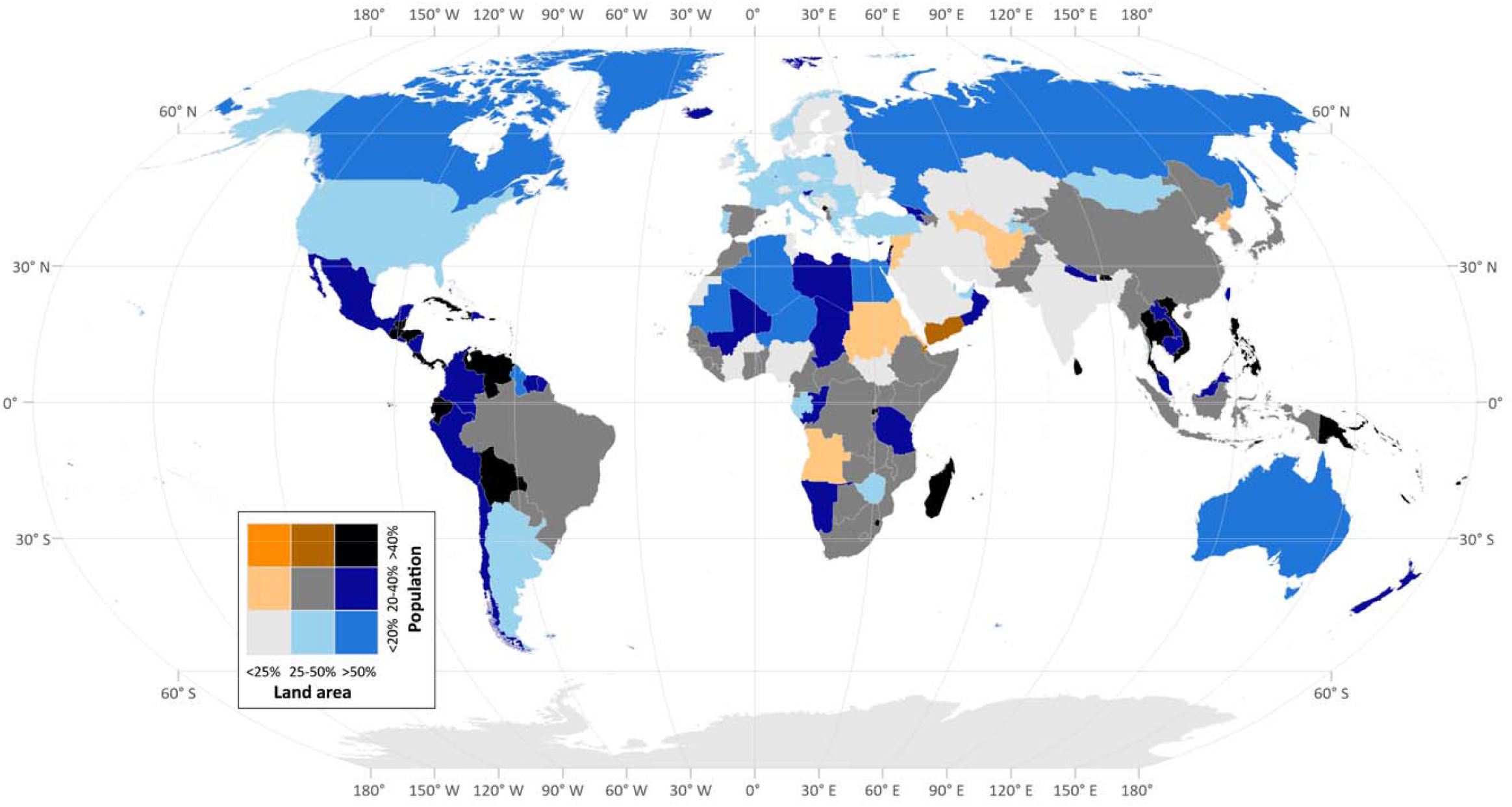
Bivariate map showing the proportion of each country’s human population living in areas requiring conservation attention, and the proportion of each country’s land area requiring conservation attention.

## Implications for global policy

Our analyses represent a comprehensive scientific estimate of the minimum land area requiring conservation attention to safeguard biodiversity. Given our inclusion of ecologically intact areas, updated maps of KBAs, and additional locations to conserve species, our estimate that 44% of land requires conservation attention is, unsurprisingly, larger than those from previous analyses that have focussed primarily on species and/or ecosystems, used earlier KBA datasets and/or didn’t include ecologically intact areas (e.g. 27.9% Butchart, et al. ^15^, 20.2% Venter, et al. ^14^, and 30% Larsen, et al. ^4^). Conservation attention to the areas we identify will be important for achieving a suite of targets in the post-2020 Global Biodiversity Framework under the Convention for Biological Diversity. These include increasing the area, connectivity and integrity of natural ecosystems, and supporting healthy and resilient populations of all species while reducing the number of species that are threatened and maintaining genetic diversity (the focus of draft Goal A); retaining ecologically intact areas (draft Target 1); conserving areas of particular importance for biodiversity (draft Target 2); and enabling recovery and conservation of wild species of fauna and flora (draft Target 3)^48^.

The figure of 44% of Earth’s land requiring conservation attention is large; however, 70% of this area is still relatively intact, implying these places may not need the larger investments required to restore landscapes^49^. In contrast, 1.3 million km^2^ of land needing conservation, mostly in developing and emerging economies, is at risk of habitat conversion to intensive human land-uses and consequent biodiversity loss so is an immediate conservation priority. Appropriately worded targets in the post-2020 Global Biodiversity Framework to safeguard these at-risk places would make a significant contribution towards addressing the biodiversity crisis, as long as it is accompanied with parallel efforts ensuring that habitat conversion is not displaced into other important conservation areas^50^.

Our finding that 1.8 billion people live in areas requiring conservation attention raises important questions about implementation. Historically, some conservation actions have adversely affected and continue to negatively affect Indigenous Peoples, Afro-descendants, and local communities^39-41^. The high number of people living in areas requiring conservation attention implies that practices such as displacing or relocating people will not only be unjust, but also not possible. Evidence shows that in many cases Indigenous Peoples and local communities have been effective stewards of biodiversity worldwide^51^. An ethical strategy that may effectively safeguard large extents of land is a rights-based approach to conservation^45,52^. The central pillars of this are i) recognising that through their customary practices Indigenous Peoples, Afro-descendants, and local communities have already demonstrated both leadership and agency in biodiversity conservation across the world^53^; ii) recognising their rights to land, benefit sharing, and institutions, and supporting efforts to strengthen these rights, so they can continue to effectively conserve their own lands; and iii) making Indigenous Peoples, Afro-descendants, and local communities partners in setting the global conservation agendas through the CBD and promoted as leaders in achieving its targets. Large areas requiring conservation attention are claimed by Indigenous Peoples, Afro-descendants, and local communities as their territories or lands so supporting them to continue conserving these places may be the most effective and efficient way to meet many biodiversity targets, while governments may need to work with them to ensure these lands are not converted to other less biodiversity friendly land-uses.

A number of additional actions are required to achieve the scale of conservation necessary to deliver positive biodiversity outcomes. On all land requiring conservation attention the expansion of roads and developments such as agriculture, forestry, and mining, needs to follow development frameworks such as the mitigation hierarchy to ensure ‘no net loss’ of biodiversity and natural ecosystems^54^. As such, mechanisms that direct developments away from important conservation areas are also crucial, including strengthening investment and performance standards for financial organisations such as the World Bank and other development investors^55^, and tightening existing industry certification standards. Our threat analysis only looked at future land conversion; however, a range of other threats such as overhunting, climate change, and fragmentation must also be considered and mitigated in areas requiring conservation attention.

A critical implementation challenge is that the proportion of land that different countries would need to conserve is highly inequitable. This variation is largely a reflection of the distribution of biodiversity, where tropical countries with high species richness and many restricted range endemics require large areas of land to be conserved because there are few other places to conserve those species. The variation is also due to the distribution of ecologically intact areas, whereby five countries, Canada, Russia, USA, Brazil and Australia contain 75% of Earth’s ecologically intact areas^17^, and so will each need to conserve large areas. In responding to this inequity, the conservation community can apply the concept of common but differentiated responsibilities that is foundational to all global environmental agendas including the CBD^56^ and United Nations Framework Convention on Climate Change^57^. Since the burden of conservation is disproportionately distributed, cost-sharing and fiscal transfer mechanisms are likely necessary to ensure that all national participation is equitable and fair, and the opportunity costs of foregone developments are considered^58,59^. This is important since the majority of land requiring conservation attention that is at risk of immediate habitat conversion is found in developing economies. Notably, many environmental impacts in emerging and developing economies are driven by consumption in developed economies^47^, who have a moral obligation to reduce these demands or fund the necessary local conservation efforts.

Our estimate of the land area requiring effective biodiversity conservation must be considered the bare minimum needed, and will almost certainly expand as more data on the distributions of underrepresented species such as plants, invertebrates, and freshwater species becomes available for future analyses^60^. New KBAs are continuing to be identified for under-represented taxonomic groups, threatened or geographically-restricted ecosystems, and highly intact and irreplaceable ecosystems. Species and ecosystems are also shifting under climate change, and as a result, are leading to changes in the location of land requiring effective conservation^61^, which we could not account for. Future analyses could use our framework to identify the efficacy of the areas we identified in conserving shifting species ranges under climate change. We also note that post-2020 biodiversity targets may imply higher levels of ecoregional representation than the 17% we used (see Methods). Many of the species representation targets (*n* = 5182, 14.6%) could not be met within existing habitat, emphasising the importance of restoration over the coming decades. Given the prioritisation approach used, every loss of a place that was identified makes the total area requiring conservation attention grow, since to meet species and ecoregion coverage targets, the algorithm will be forced to find a less-optimal configuration of land areas.

For the above reasons, our results do not imply that the land our analysis did not identify, (i.e. the other 56% of Earth’s land surface), is unimportant for conservation and global sustainable development goals. Much of this area will be important for sustaining the provision of ecosystem services to people, from climate regulation to provisioning of food, materials, drinking water, and crop pollination, in addition to supporting other elements of biodiversity not captured in our priority areas^6^. Furthermore, many human activities can impact the entire Earth system regardless of where they occur (e.g. fossil fuel use, pesticide use, and pollution), so management efforts focussed on limiting the ultimate drivers of biodiversity loss are essential^62^. Finally, we have not considered how constraining developments to locations outside of the land area needing conservation impacts solutions for meeting human needs, such as increasing energy and food demands. Although social objectives that benefit humanity are clearly important, they cannot all be achieved sustainably without limiting the degradation of the ecosystems supporting all life^1^. Integrated assessments of how we can achieve multiple social objectives while effectively conserving biodiversity at a global scale are important avenues for future research^63^.

The world’s nations are discussing post-2020 biodiversity conservation targets within the CBD and wider Sustainable Development Goals international agenda. These targets will define the global conservation agenda for at least the next decade, so it is crucial that they are adequate to achieve biodiversity outcomes^10^. Our analyses show that a minimum of 44% of land requires conservation attention, through both site- and landscape-scale approaches, which should serve as an ecological foundation for negotiations. Governments failed to meet the CBDs previous Aichi Targets suggesting a need to reimagine how conservation is done^64^. Our finding that over 1.8 billion people live on lands requiring conservation attention further supports the need for dramatic shifts in conservation strategies. The implementation of conservation actions must put the rights of Indigenous Peoples and local communities, socio-environmental justice and human rights frameworks at their centre. As such, conservation scientists have an opportunity to scale up their role as capacity builders for the communities that request their expertise. If CBD signatory nations are serious about safeguarding the biodiversity and ecosystem services that underpin all life on earth^1,63^, then they need to recognise that conservation action must be immediately and substantially scaled-up, in extent, intensity, sophistication and effectiveness.

## Methods

### Mapping important conservation areas

We obtained spatial data on the location of protected areas from the February 2020 version of the World Database on Protected Areas (WDPA)^27^. This edition does not contain data on protected areas in China, which have largely been removed from the publicly accessible WDPA in more recent versions. We therefore used the January 2017 version of the WDPA for China, since this is the most recent version with China’s full complement of protected areas. In total, we had location data for 253,797 protected areas. We handled the WDPA data according to best-practice guidelines that are available on the protected planet website (https://www.protectedplanet.net/c/calculating-protected-areacoverage) and included regionally, nationally and internationally designated protected areas. We included all protected areas in the database regardless of their IUCN management category because these categories are not globally consistent. The WDPA dataset contains protected areas represented as point data. In these cases, we converted the points to polygons by setting a geodesic buffer around the point based on the areal attributes of that point. We excluded points with no areal attributes. We also excluded all marine protected areas, ‘proposed’ protected areas, and UNESCO Man and Biosphere Reserves since their core conservation areas often overlap with other protected areas and their buffer zones’ primary goals are not biodiversity conservation. Finally, we flattened (i.e. dissolved) the protected area data to remove any overlapping protected areas.

We obtained data on the boundaries of 14,192 KBAs from the September 2019 version of the World Database of Key Biodiversity Areas^28^. KBAs documented with point data were treated as outlined above for protected areas. The KBA dataset includes sites identified under previously established criteria such as Important Bird and Biodiversity Areas (IBAs)^65^ and Alliance for Zero Extinction sites (AZEs)^66^ (as the KBA Standard explicitly state that these sites are encompassed by KBAs), and the KBA criteria builds closely on these previous criteria^16^. Although the KBA criteria have been applied most comprehensively to birds, in the September 2019 version of the KBA dataset, 53% of species that trigger the criteria are non-avian, and 35% of sites are triggered by non-avian species. These proportions are increasing as the standard is applied more widely to non-birds, and many bird-triggered KBAs are likely to prove important for other species^65^. We obtained global data on the extent of ecologically intact areas from Allan, et al. ^29^, who utilised maps of ‘pressure-free lands’. Previous analyses have referred to these pressure free lands as wilderness areas, but here we avoid the term, preferring ‘ecologically intact’ since the word wilderness is sometimes associated with a legacy of violence that has been perpetrated to promote it and is therefore offensive to some people.

We merged protected areas, KBAs and ecologically intact areas together, removing overlaps (i.e. again flattened the merged datasets) to create a global template of “existing important conservation areas”.

### Distribution and representation of biodiversity

We obtained data on the distributions of terrestrial mammals (*n* = 5,617), amphibians (*n* = 6,577), freshwater crabs (*n* = 1,285), shrimp (*n* = 692) and crayfish (*n* = 496) from the IUCN Red List of Threatened Species^67^. Bird distribution data (*n* = 10,926) were sourced from BirdLife International and Handbook of the Birds of the World^68^, and reptile data (*n* = 9,964) from Roll, et al. ^69^. These represent the most comprehensive spatial databases for these taxonomic groups. We excluded species that are extinct, possibly extinct, or if their presence is uncertain. We did not account for sub-species. The freshwater species ranges are mapped at the watershed level which is generally coarser than the 30 × 30 km resolution of our spatial analysis. Since freshwater species are likely to only inhabit a small area within the watersheds, there is a chance of commission errors, where a species is falsely identified as present. In regions with larger hydro sheds the probability of commission errors increases. There is also a higher likelihood of commission errors in less surveyed regions such as the global tropics, where there are also many narrow-ranged species. This is important information for interpreting the results, and highlights the need for downscaled national level analyses using best available local data. We also included data on the distribution of 845 terrestrial ecoregions^31^, which are bio-geographically distinct spatial units at the global scale.

We set representation targets for the percentage of each species’ distribution that should be effectively conserved, following previous studies (Rodrigues, et al. ^30^, Venter, et al. ^14^, and Butchart, et al. ^15^). Targets were set as a function of a species’ range size, and were log-linearly scaled between 10% for species with distributions >250,000 km^2^, to 100% for species with ranges <1,000 km^2^. We limited the target for species with large ranges to 1 million km^2^ maximum^15^. We acknowledge that other target setting approaches exist, for example based on minimising species extinction risk^70^. However, these are not as widely adopted as the approach we followed here. We also acknowledge that scaling targets for species based on range size may not always be sufficient to guarantee persistence for all species. That said, it is the most widely used “best practice” target setting approach. For each ecoregion, we followed Venter et al. ^14^ by setting a coverage target of 17%, in line with Aichi Target 11 of the Strategic Plan for Biodiversity^3^. We acknowledge that Aichi Target 11 expired in 2020 but the nature of the post-2020 targets is still under discussion, and that the 17% value is arbitrary and was determined through negotiation. We carried out a “gap analysis” by calculating the proportion of each species’ range that currently overlaps with the important conservation areas, and comparing this with each species’ coverage target to identify under-represented species and the extent of additional range each requires.

### Priority areas for the expansion of conservation efforts

We identify spatial priorities for meeting species conservation targets, whilst accounting for current protection within existing important conservation areas, and minimizing the cost (the area of a planning unit) of the areas selected^71^. We solve this using the mathematical optimisation ‘minimum set problem’ (also known as the ‘reserve selection problem’), an integer linear programming problem, using Gurobi (version 5.6.2) following the methods developed by Beyer, et al. ^72^. Integer linear programming is an effective, exact method for solving optimisation problems, which minimises or maximises an objective function subject to constraints conditional on the decision variables being integers^72^. Specifically, we solved the reserve selection problem as follows:

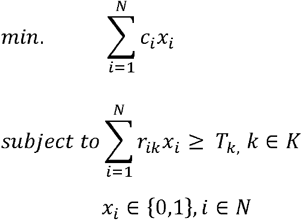

where *x*_*i*_ is a binary decision variable determining whether planning unit *i* is selected (1) or not (0), and *c*_*i*_ represents the cost of planning unit *i* or, in this case the objective is to select the smallest number of planning units, so *c*_*i*_ = 1 for every *i*. The parameter *r*_*ik*_ is the contribution of planning unit *i* to feature *k* and *T*_*k*_ is the minimum target (described above) to be achieved for feature *k* among all planning units. We applied a threshold specifying that solutions must be within 0.5% of the optimum^72^, which returns a near-optimal solution.

To run the analysis, we first created a 30 × 30 km (900 km^2^) global planning unit grid. This resolution limits the risk of commission errors when working with the available species distribution data (e.g. assuming a species is present when it is not)^15,73^. Planning units were clipped to terrestrial areas and inland lakes and waterways so that freshwater taxa could be included. We included Antarctica and Greenland. We calculated the area of each conservation feature (e.g. species distribution and ecoregion distribution) within each planning unit, including the area within existing important conservation areas. All geospatial data processing was carried out in the Mollweide equal-area projection using a spatially enabled PostgreSQL database (using PostGIS version 2.2) or in ESRI ArcGIS version 10.5.1.

We used the area of a planning unit as a surrogate for the cost of conservation in that planning unit. Seeking to minimise area is advantageous because it supports our aim of identifying the minimum area requiring conservation attention globally. There is also evidence that area is a good proxy for cost, reducing uncertainties created in the absence of fine scale and accurate cost data^74^. Other widely used cost metrics such as the human footprint^75,76^ and agricultural opportunity^77^ costs do not extend to Antarctica or remote sub-Antarctic islands further supporting our choice of area as the most suitable cost metric.

To explore how sensitive our results are to the choice of cost metric we ran the prioritisation analyses again using two other cost layers: the sum human footprint and the agricultural opportunity cost of a planning unit. The human footprint is a map of cumulative human pressure on the natural environment for the year 2009 at a 1km^2^ resolution globally^75,76^. The agricultural opportunity data is a global map of the gross economic rents of agricultural lands^77^. We assumed that conservation will be cheaper and more feasible in areas with less human influence and lower agricultural opportunity. We excluded Antarctica and sub-Antarctic islands from this sensitivity analysis. We found that the priorities identified using different cost layers overlap by 58% on average (ranging from 36–75% overlap) (Extended Data Table 7; Extended Data Figure 3). This demonstrates that our results are somewhat sensitive to cost, but are also driven to large extent by the distribution of biodiversity features.

We accounted for current land-use in our analyses by excluding places classified as ‘built areas’, assuming they are unavailable for conservation. By built areas we mean cities and major urban centres that contain no original habitat. Data on the extent of built areas was obtained from the European Space Agency (ESA) Climate Change Initiative (CCI) who have developed globally consistent landcover maps at a 300 m resolution for the year 2015, classing the world into 22 land use categories^78^. We extracted land use category 190 which represents urban areas and resampled the data to a 1 km^2^ resolution were a pixel was considered a built area if >50% of its area was urban. In the results presented in the main manuscript we assumed all other land-uses including current agricultural areas are available for conservation since they can be restored, and our aim is to identify the ‘minimum area requiring conservation attention’ even if that means including places requiring restoration. Some KBAs contain urban areas because the management units they represent contain such urban areas, or, more rarely, they support significant populations of species of conservation concern in these locations. We did not account for this in the analyses, so the urban extent of these KBAs would have been considered unavailable for meeting species representation targets. This means that the 44% of Earth’s surface that we calculated is a slight underestimate of the true extent requiring conservation attention.

To assess the sensitivity of our results to current land use we ran the prioritisation again excluding both built areas and current agricultural extent, assuming this land is unavailable for conservation (Extended Data Figure 3). Data on agricultural extent was also obtained from the ESA CCI^78^. We extracted land-use categories 10; rainfed cropland, 11; herbaceous cover, 12; tree or shrub cover, 20; irrigated or post flooding cropland, and 30; mosaic cropland, converted this into a binary agriculture is present/absent layer and resampled to 1 km^2^ resolution where a pixel was considered agriculture if >50% of its area was covered by agricultural land-use. We found that when we exclude both built areas and agricultural land (and used planning unit area as a cost) the land area requiring conservation is 695,633 km^2^ lower than when agriculture was included. However, this is because 5,182 species (14.6%) cannot meet their representation targets when the model cannot select areas under current agriculture, resulting in an insufficient conservation plan.

By running the prioritisation with different cost layers and land-use constraints, we identify different spatial solutions that meet the species distribution coverage targets. This demonstrates that there is considerable spatial freedom in identifying priority conservation areas. The fact that not all targets could be met when agricultural and urban land was locked out also demonstrates the bounds of this freedom. Finding multiple near optimal spatial solutions to conservation planning problems is one of the most important functionalities of conservation planning tools since it allows decision makers to assess multiple options for achieving their goals.

It is possible to create conservation plans where each country must conserve the same proportion of their area^79^; however, this leads to costly inefficient plans^59^, and would be inconsistent with our aim of identifying the minimum most important area requiring conservation. Therefore, we ran the prioritisation at the global scale.

### Future threats to conservation areas

To map the risk of habitat conversion occurring in the conservation areas identified, we utilised spatially explicit data on future land-use scenarios from the newly released Land Use Harmonisation Dataset v2 (http://luh.umd.edu/)^35^. To determine optimistic, middle-of-the-road, and pessimistic scenarios, we evaluated projections under three different Shared Socioeconomic Pathways (SSPs)^36^, which are linked to Representative Concentration Pathways (RCPs)^37^: specifically, SSP1 (RCP2.6; IMAGE), an optimistic scenario where the world gradually moves towards a more sustainable future, SSP2 (MESSAGE-GLOBIOM) a middle-of-the-road scenario without any extreme changes towards or away from sustainability, and SSP3 (RCP7.0; AIM), a pessimistic scenario where land use change is poorly regulated.

The harmonised land-use data contains 12 state layers (with the unit being the fraction of a grid cell in that state) for the years 2015 (current baseline), 2030 and 2050. We considered four of the state layers as natural land-cover classes, including; primary forested land, primary non-forested land, potentially forested secondary land, and potentially non-forested secondary land (Extended Data Figure 4). Using these four classes, we calculated the proportion of natural land projected to be lost (converted to human uses) by the years 2030 and 2050 in each 30 x 30 km grid cell. From this we calculated the area of natural land projected to be lost within each grid cell. We assume that once land is converted it remains converted. Antarctica and remote islands were excluded from this part of the analyses because the land-use data does not extend to them. We also created an “ensemble” scenario, where we calculated the average area of natural land projected to be converted in each pixel across all three SSPs (Extended Data Figure 5).

### Estimating the human population in areas requiring conservation

We used LandScan’s global population distribution model for the year 2018^38^ to estimate the number of people living within areas requiring conservation. We expanded on methods used by Schleicher et al.^80^, who used LandScan to measure the populations living in the least populated ecoregions. Data were extracted to estimate the area and number of people found within places requiring conservation. These were then tabulated using the database of Global Administrative Areas (GADM 2020) to provide measures for each territory. Population data were calculated in raster format at a resolution of 30 by 30 arc seconds, approximately 1 km^2^ (835m^2^). LandScan population data represents an ambient population (average over 24 hours).

## Acknowledgements

We thank Peadar Brehony, Peter Tyrell, Chris Sandbrook, Oscar Venter and Piero Visconti for thoughtful comments on the manuscript.

## Author Contributions

J.R.A, J.E.M.W, and H.P.P. framed the study. J.R.A., S.C.A., M.D.M., G.G., P.M. carried out the analyses. All authors discussed and interpreted the results. J.R.A wrote the manuscript with support from all authors.

