## Extended Data for "The minimum land area requiring conservation attention to safeguard biodiversity"

**Abbreviated Title:** Land area needed to conserve biodiversity

**Authors**

James R. Allan*^1,2^, Hugh P. Possingham^2,3^, Scott C. Atkinson^2,4^, Anthony Waldron^5^, Moreno Di Marco^6,7^, Vanessa M. Adams^8^, Stuart H. M. Butchart^9,10^, W. Daniel Kissling^1^, Thomas Worsdell^11^, Gwili Gibbon^12^, Kundan Kumar^11^, Piyush Mehta^13^, Martine Maron^2,7^, Brooke A. Williams^2,7^, Kendall R. Jones^14^, Brendan A. Wintle^15^, April E. Reside^2,7^, James E.M. Watson^2,7^

**Affiliations**

^1^Institute for Biodiversity and Ecosystem Dynamics (IBED), University of Amsterdam, P.O. Box 94240, 1090 GE, Amsterdam, The Netherlands

^2^Centre for Biodiversity and Conservation Science, The University of Queensland, St Lucia, QLD 4072, Australia

^3^The Nature Conservancy, VA 22203-1606, USA

^4^United Nations Development Programme (UNDP), New York, New York, USA

^5^Cambridge Conservation Initiative, David Attenborough Building, Department of Zoology, Cambridge University, Cambridge CB2 3QZ, UK

^6^Department of Biology and Biotechnologies, Sapienza University of Rome, viale dell'Università 32, I-00185 Rome, Italy

^7^School of Earth and Environmental Sciences, The University of Queensland, St Lucia QLD 4072, Australia

^8^School of Technology, Environments & Design, University of Tasmania, Hobart, TAS 7001, Australia

^9^BirdLife International, David Attenborough Building, Pembroke Street, Cambridge CB2 3QZ, UK

^10^Department of Zoology, University of Cambridge, Downing Street, Cambridge CB2 3EJ, UK

^11^Rights and Resources Initiative, Washington, D. C., USA

^12^Durrell Institute of Conservation and Ecology, School of Anthropology and Conservation, University of Kent, Canterbury, CT2 7NR, UK

^13^University of Delaware, Newark, DE 19716, USA

^14^Wildlife Conservation Society, Global Conservation Program, 2300 Southern Boulevard, Bronx, NY 10460-1068, USA

^15^School of BioSciences, University of Melbourne, Vic., Australia

**Keywords** Protected Areas, Conservation Planning, Restoration, Global Conservation Priorities, Strategic Plan for Biodiversity, Convention on Biological Diversity, Conservation Priorities, Key Biodiversity Areas, Wilderness, Ecologically Intact.

| ***Extended Data Table 1****. The area and proportion of each country identified as important for conservation* | | | | |
| --- | --- | --- | --- | --- |
| **Country** | **Continent** | **Country area (km)** | **Conservation area (km)** | **Conservation percent (%)** |
| Afghanistan | Asia | 643831 | 132899 | 20.6 |
| Akrotiri | Asia | 99 | 86 | 87.4 |
| Aland Islands | Europe | 939 | 12 | 1.3 |
| Albania | Europe | 28362 | 11508 | 40.6 |
| Algeria | Africa | 2317479 | 1293468 | 55.8 |
| American Samoa | Oceania | 181 | 54 | 29.6 |
| Andorra | Europe | 453 | 442 | 97.7 |
| Angola | Africa | 1252237 | 270477 | 21.6 |
| Anguilla | North America | 81 | 80 | 99.1 |
| Antarctica | Antarctica | 12078023 | 26954 | 0.2 |
| Antigua and Barbuda | North America | 454 | 443 | 97.7 |
| Argentina | South America | 2790757 | 984057 | 35.3 |
| Armenia | Asia | 29621 | 17563 | 59.3 |
| Aruba | North America | 171 | 159 | 92.9 |
| Ashmore and Cartier Islands | Oceania | 3 | 0 | 0.0 |
| Australia | Oceania | 7723183 | 4019833 | 52.0 |
| Austria | Europe | 83943 | 28141 | 33.5 |
| Azerbaijan | Asia | 86346 | 31743 | 36.8 |
| Bahamas | North America | 12649 | 11254 | 89.0 |
| Bahrain | Asia | 587 | 245 | 41.7 |
| Baikonur Cosmodrome | Asia | 6500 | 0 | 0.0 |
| Bangladesh | Asia | 137524 | 34567 | 25.1 |
| Barbados | North America | 447 | 397 | 88.8 |
| Belarus | Europe | 207095 | 35450 | 17.1 |
| Belgium | Europe | 30630 | 9608 | 31.4 |
| Belize | North America | 22421 | 16548 | 73.8 |
| Benin | Africa | 116851 | 42605 | 36.5 |
| Bermuda | North America | 61 | 59 | 97.1 |
| Bhutan | Asia | 40522 | 26021 | 64.2 |
| Bolivia | South America | 1092895 | 566673 | 51.9 |
| Bosnia and Herzegovina | Europe | 51837 | 9204 | 17.8 |
| Botswana | Africa | 581815 | 252839 | 43.5 |
| Brazil | South America | 8523997 | 4206016 | 49.3 |
| British Indian Ocean Territory | Oceanic Islands | 49 | 49 | 100.0 |
| British Virgin Islands | North America | 146 | 133 | 91.1 |
| Brunei Darussalam | Asia | 5755 | 3785 | 65.8 |
| Bulgaria | Europe | 112820 | 46724 | 41.4 |
| Burkina Faso | Africa | 274440 | 44055 | 16.1 |
| Burundi | Africa | 27222 | 8756 | 32.2 |
| Cambodia | Asia | 182161 | 92085 | 50.6 |
| Cameroon | Africa | 467376 | 165963 | 35.5 |
| Canada | North America | 9916542 | 8306743 | 83.8 |
| Cape Verde | Africa | 3905 | 3890 | 99.6 |
| Cayman Islands | North America | 311 | 301 | 96.7 |
| Central African Republic | Africa | 622033 | 249602 | 40.1 |
| Chad | Africa | 1273567 | 702961 | 55.2 |
| Chile | South America | 738018 | 427446 | 57.9 |
| China | Asia | 9393754 | 3527688 | 37.6 |
| Clipperton Island | Oceanic Islands | 5 | 5 | 100.0 |
| Colombia | South America | 1142648 | 722797 | 63.3 |
| Comoros | Africa | 1682 | 1673 | 99.4 |
| Cook Islands | Oceania | 192 | 187 | 97.3 |
| Coral Sea Islands | Oceania | 0 | 0 | 99.3 |
| Costa Rica | North America | 51469 | 44007 | 85.5 |
| Cote d'Ivoire | Africa | 322760 | 91748 | 28.4 |
| Croatia | Europe | 55078 | 25278 | 45.9 |
| Cuba | North America | 110469 | 73485 | 66.5 |
| Curacao | North America | 466 | 405 | 87.0 |
| Cyprus | Asia | 5408 | 3786 | 70.0 |
| Cyprus U.N. Buffer Zone | Asia | 342 | 135 | 39.5 |
| Czech Republic | Europe | 78673 | 17502 | 22.2 |
| Dem. Rep. Korea | Asia | 122520 | 24970 | 20.4 |
| Democratic Republic of the Congo | Africa | 2340650 | 630995 | 27.0 |
| Denmark | Europe | 42590 | 10061 | 23.6 |
| Dhekelia | Asia | 128 | 26 | 20.2 |
| Djibouti | Africa | 21982 | 7654 | 34.8 |
| Dominica | North America | 735 | 733 | 99.7 |
| Dominican Republic | North America | 48698 | 41324 | 84.9 |
| Ecuador | South America | 256729 | 209067 | 81.4 |
| Egypt | Africa | 1005139 | 615469 | 61.2 |
| El Salvador | North America | 20660 | 12470 | 60.4 |
| Equatorial Guinea | Africa | 26851 | 7592 | 28.3 |
| Eritrea | Africa | 123248 | 30355 | 24.6 |
| Estonia | Europe | 45678 | 9668 | 21.2 |
| Ethiopia | Africa | 1134603 | 423168 | 37.3 |
| Faeroe Islands | Europe | 1324 | 1088 | 82.2 |
| Falkland Islands | South America | 11579 | 7176 | 62.0 |
| Federated States of Micronesia | Oceania | 638 | 601 | 94.2 |
| Fiji | Oceania | 19034 | 16245 | 85.3 |
| Finland | Europe | 331677 | 79013 | 23.8 |
| France | Europe | 637161 | 229560 | 36.0 |
| French Polynesia | Oceania | 3356 | 3339 | 99.5 |
| French Southern and Antarctic Lands | Oceanic Islands | 7273 | 7269 | 99.9 |
| Gabon | Africa | 261718 | 85140 | 32.5 |
| Georgia | Asia | 69618 | 35655 | 51.2 |
| Germany | Europe | 357177 | 149887 | 42.0 |
| Ghana | Africa | 240213 | 78903 | 32.8 |
| Gibraltar | Europe | 4 | 4 | 97.9 |
| Greece | Europe | 131538 | 64540 | 49.1 |
| Greenland | North America | 2142675 | 2111379 | 98.5 |
| Grenada | North America | 349 | 346 | 99.1 |
| Guam | Oceania | 568 | 568 | 100.0 |
| Guatemala | North America | 109439 | 86870 | 79.4 |
| Guernsey | Europe | 73 | 8 | 10.7 |
| Guinea | Africa | 245839 | 122007 | 49.6 |
| Guinea-Bissau | Africa | 33031 | 14986 | 45.4 |
| Guyana | South America | 212614 | 113397 | 53.3 |
| Haiti | North America | 27035 | 25312 | 93.6 |
| Heard I. and McDonald Islands | Oceanic Islands | 394 | 394 | 100.0 |
| Honduras | North America | 112894 | 73659 | 65.2 |
| Hong Kong | Asia | 1041 | 902 | 86.6 |
| Hungary | Europe | 93155 | 23422 | 25.1 |
| Iceland | Europe | 101950 | 53823 | 52.8 |
| India | Asia | 3166098 | 789331 | 24.9 |
| Indian Ocean Territories | Asia | 113 | 113 | 100.0 |
| Indonesia | Asia | 1892367 | 915861 | 48.4 |
| Iran | Asia | 1627124 | 378343 | 23.3 |
| Iraq | Asia | 438564 | 69000 | 15.7 |
| Ireland | Europe | 69315 | 12248 | 17.7 |
| Isle of Man | Europe | 574 | 197 | 34.3 |
| Israel | Asia | 21968 | 17521 | 79.8 |
| Italy | Europe | 301351 | 123631 | 41.0 |
| Jamaica | North America | 11093 | 10818 | 97.5 |
| Japan | Asia | 374168 | 162418 | 43.4 |
| Jersey | Europe | 119 | 19 | 16.4 |
| Jordan | Asia | 89140 | 19634 | 22.0 |
| Kazakhstan | Asia | 2712413 | 447379 | 16.5 |
| Kenya | Africa | 589637 | 196698 | 33.4 |
| Kiribati | Oceania | 956 | 850 | 88.8 |
| Kosovo | Europe | 10919 | 3491 | 32.0 |
| Kuwait | Asia | 17535 | 4151 | 23.7 |
| Kyrgyzstan | Asia | 199157 | 53083 | 26.7 |
| Lao PDR | Asia | 229339 | 114445 | 49.9 |
| Latvia | Europe | 64403 | 11762 | 18.3 |
| Lebanon | Asia | 10025 | 7016 | 70.0 |
| Lesotho | Africa | 30211 | 12451 | 41.2 |
| Liberia | Africa | 95924 | 45555 | 47.5 |
| Libya | Africa | 1630170 | 994319 | 61.0 |
| Liechtenstein | Europe | 137 | 48 | 34.9 |
| Lithuania | Europe | 64786 | 12407 | 19.2 |
| Luxembourg | Europe | 2606 | 1349 | 51.8 |
| Macao | Asia | 30 | 21 | 69.4 |
| Macedonia | Europe | 25406 | 11015 | 43.4 |
| Madagascar | Africa | 596083 | 315353 | 52.9 |
| Malawi | Africa | 120117 | 56876 | 47.4 |
| Malaysia | Asia | 330072 | 205980 | 62.4 |
| Maldives | Oceanic Islands | 109 | 107 | 97.7 |
| Mali | Africa | 1259626 | 665267 | 52.8 |
| Malta | Europe | 326 | 271 | 82.9 |
| Marshall Islands | Oceania | 166 | 98 | 59.2 |
| Mauritania | Africa | 1041681 | 683533 | 65.6 |
| Mauritius | Oceanic Islands | 2025 | 1939 | 95.7 |
| Mexico | North America | 1966640 | 988604 | 50.3 |
| Moldova | Europe | 33190 | 2365 | 7.1 |
| Monaco | Europe | 19 | 15 | 78.7 |
| Mongolia | Asia | 1564041 | 477977 | 30.6 |
| Montenegro | Europe | 13735 | 7084 | 51.6 |
| Montserrat | North America | 100 | 100 | 100.0 |
| Morocco | Africa | 593757 | 221737 | 37.3 |
| Mozambique | Africa | 792793 | 253469 | 32.0 |
| Myanmar | Asia | 666446 | 248153 | 37.2 |
| Namibia | Africa | 826671 | 423939 | 51.3 |
| Nauru | Oceania | 29 | 29 | 100.0 |
| Nepal | Asia | 147653 | 73313 | 49.7 |
| Netherlands | Europe | 37337 | 13687 | 36.7 |
| New Caledonia | Oceania | 18934 | 18683 | 98.7 |
| New Zealand | Oceania | 268706 | 155131 | 57.7 |
| Nicaragua | North America | 129473 | 69505 | 53.7 |
| Niger | Africa | 1187814 | 695032 | 58.5 |
| Nigeria | Africa | 913257 | 210828 | 23.1 |
| Niue | Oceania | 222 | 41 | 18.4 |
| Norfolk Island | Oceania | 41 | 41 | 100.0 |
| Northern Cyprus | Asia | 3327 | 1803 | 54.2 |
| Northern Mariana Islands | Oceania | 583 | 582 | 99.9 |
| Norway | Europe | 380388 | 165285 | 43.5 |
| Oman | Asia | 312789 | 157643 | 50.4 |
| Pakistan | Asia | 875892 | 229294 | 26.2 |
| Palau | Oceania | 482 | 481 | 99.8 |
| Palestine | Asia | 6298 | 4632 | 73.5 |
| Panama | North America | 75011 | 59982 | 80.0 |
| Papua New Guinea | Oceania | 468205 | 266076 | 56.8 |
| Paraguay | South America | 401756 | 155822 | 38.8 |
| Peru | South America | 1298027 | 801709 | 61.8 |
| Philippines | Asia | 295037 | 190770 | 64.7 |
| Pitcairn Islands | Oceania | 43 | 43 | 99.6 |
| Poland | Europe | 312917 | 132449 | 42.3 |
| Portugal | Europe | 91507 | 26953 | 29.5 |
| Puerto Rico | North America | 9055 | 8066 | 89.1 |
| Qatar | Asia | 11198 | 4016 | 35.9 |
| Republic of Congo | Africa | 347204 | 178575 | 51.4 |
| Republic of Korea | Asia | 98741 | 41748 | 42.3 |
| Romania | Europe | 236334 | 77165 | 32.7 |
| Russian Federation | Asia | 16896872 | 10487173 | 62.1 |
| Rwanda | Africa | 25475 | 13000 | 51.0 |
| Saint Helena | Oceanic Islands | 368 | 357 | 97.0 |
| Saint Kitts and Nevis | North America | 266 | 261 | 98.3 |
| Saint Lucia | North America | 609 | 602 | 98.9 |
| Saint Pierre and Miquelon | North America | 243 | 23 | 9.5 |
| Saint Vincent and the Grenadines | North America | 370 | 367 | 99.1 |
| Saint-Bartholemy | North America | 25 | 25 | 99.9 |
| Saint-Martin | North America | 69 | 66 | 96.0 |
| Samoa | Oceania | 2797 | 2433 | 87.0 |
| San Marino | Europe | 60 | 0 | 0.0 |
| Sao Tome and Principe | Africa | 1044 | 1030 | 98.7 |
| Saudi Arabia | Asia | 1930318 | 249670 | 12.9 |
| Scarborough Reef | Asia | 0 | 0 | 0.0 |
| Senegal | Africa | 197383 | 75556 | 38.3 |
| Serbia | Europe | 77589 | 12246 | 15.8 |
| Serranilla Bank | North America | 0 | 0 | 0.0 |
| Seychelles | Oceanic Islands | 411 | 394 | 95.9 |
| Siachen Glacier | Asia | 2092 | 938 | 44.8 |
| Sierra Leone | Africa | 72073 | 22120 | 30.7 |
| Singapore | Asia | 514 | 255 | 49.6 |
| Sint Maarten | North America | 23 | 20 | 85.3 |
| Slovakia | Europe | 48417 | 19204 | 39.7 |
| Slovenia | Europe | 20322 | 11475 | 56.5 |
| Solomon Islands | Oceania | 27373 | 22939 | 83.8 |
| Somalia | Africa | 474931 | 134029 | 28.2 |
| Somaliland | Africa | 168470 | 51811 | 30.8 |
| South Africa | Africa | 1224180 | 420867 | 34.4 |
| South Georgia and South Sandwich Islands | Oceanic Islands | 3913 | 3913 | 100.0 |
| South Sudan | Africa | 630939 | 150726 | 23.9 |
| Spain | Europe | 507461 | 252683 | 49.8 |
| Spratly Islands | Asia | 12 | 0 | 0.0 |
| Sri Lanka | Asia | 66722 | 51067 | 76.5 |
| Sudan | Africa | 1868197 | 308247 | 16.5 |
| Suriname | South America | 146091 | 123175 | 84.3 |
| Swaziland | Africa | 17181 | 10333 | 60.1 |
| Sweden | Europe | 444490 | 103230 | 23.2 |
| Switzerland | Europe | 41419 | 8960 | 21.6 |
| Syria | Asia | 186367 | 35270 | 18.9 |
| Taiwan | Asia | 36362 | 29345 | 80.7 |
| Tajikistan | Asia | 142454 | 69957 | 49.1 |
| Tanzania | Africa | 947673 | 521492 | 55.0 |
| Thailand | Asia | 517430 | 261326 | 50.5 |
| The Gambia | Africa | 10564 | 8720 | 82.5 |
| Timor-Leste | Asia | 15179 | 8508 | 56.0 |
| Togo | Africa | 57229 | 18793 | 32.8 |
| Tonga | Oceania | 606 | 432 | 71.2 |
| Trinidad and Tobago | North America | 5155 | 4538 | 88.0 |
| Tunisia | Africa | 157005 | 27532 | 17.5 |
| Turkey | Asia | 781171 | 251554 | 32.2 |
| Turkmenistan | Asia | 471503 | 90016 | 19.1 |
| Turks and Caicos Islands | North America | 447 | 444 | 99.4 |
| Tuvalu | Oceania | 23 | 2 | 10.5 |
| Uganda | Africa | 243480 | 86798 | 35.6 |
| Ukraine | Europe | 598550 | 97920 | 16.4 |
| United Arab Emirates | Asia | 71407 | 24530 | 34.4 |
| United Kingdom | Europe | 243280 | 82811 | 34.0 |
| United States | North America | 9469111 | 3562220 | 37.6 |
| United States Minor Outlying Islands | North America | 25 | 25 | 100.0 |
| United States Virgin Islands | North America | 357 | 299 | 83.7 |
| Uruguay | South America | 177840 | 51604 | 29.0 |
| US Naval Base Guantanamo Bay | North America | 66 | 66 | 100.0 |
| Uzbekistan | Asia | 448182 | 106763 | 23.8 |
| Vanuatu | Oceania | 12367 | 9156 | 74.0 |
| Venezuela | South America | 918623 | 651611 | 70.9 |
| Vietnam | Asia | 330727 | 206180 | 62.3 |
| Wallis and Futuna Islands | Oceania | 140 | 24 | 16.9 |
| Western Sahara | Africa | 90900 | 24180 | 26.6 |
| Yemen | Asia | 455667 | 157598 | 34.6 |
| Zambia | Africa | 756420 | 350588 | 46.3 |
| Zimbabwe | Africa | 391404 | 119839 | 30.6 |

| ***Extended Data Table 2.*** *The area and proportion of each country that is currently a protected area (PA), Key Biodiversity Area (KBA), Ecologically intact area (EIA), the combination of the three excluding their overlaps, and the newly identified additional priority areas.* | | | | | | | | | | |
| --- | --- | --- | --- | --- | --- | --- | --- | --- | --- | --- |
|  | **PA** | | **KBA** | | **EIA** | | **PA + KBA + EIA** | | **New Priorities** | |
| **Country** | **Area (km)** | **Percent (%)** | **Area (km)** | **Percent (%)** | **Area (km)** | **Percent (%)** | **Area (km)** | **Percent (%)** | **Area (km)** | **Percent (%)** |
| Afghanistan | 1381 | 0.2 | 69662 | 10.8 | 7072 | 1.1 | 70394 | 10.9 | 62506 | 9.7 |
| Akrotiri | 75 | 76.0 | 60 | 61.0 | 0 | 0.0 | 84 | 84.9 | 2 | 2.5 |
| Aland Islands | 7 | 0.7 | 0 | 0.0 | 0 | 0.0 | 7 | 0.7 | 5 | 0.6 |
| Albania | 5269 | 18.6 | 6234 | 22.0 | 0 | 0.0 | 8592 | 30.3 | 2916 | 10.3 |
| Algeria | 175730 | 7.6 | 199725 | 8.6 | 1137865 | 49.1 | 1261850 | 54.4 | 31618 | 1.4 |
| American Samoa | 29 | 16.2 | 39 | 21.6 | 0 | 0.0 | 43 | 23.5 | 11 | 6.0 |
| Andorra | 140 | 30.9 | 440 | 97.2 | 0 | 0.0 | 442 | 97.7 | 0 | 0.0 |
| Angola | 86954 | 6.9 | 78861 | 6.3 | 0 | 0.0 | 105297 | 8.4 | 165180 | 13.2 |
| Anguilla | 11 | 13.7 | 17 | 21.5 | 0 | 0.0 | 29 | 35.1 | 52 | 64.0 |
| Antarctica | 0 | 0.0 | 466 | 0.0 | 0 | 0.0 | 26954 | 0.2 | 0 | 0.0 |
| Antigua and Barbuda | 94 | 20.8 | 91 | 20.0 | 0 | 0.0 | 128 | 28.3 | 315 | 69.4 |
| Argentina | 235500 | 8.4 | 297184 | 10.6 | 141219 | 5.1 | 508186 | 18.2 | 475871 | 17.1 |
| Armenia | 7407 | 25.0 | 10084 | 34.0 | 616 | 2.1 | 13227 | 44.7 | 4336 | 14.6 |
| Aruba | 32 | 18.7 | 37 | 21.4 | 0 | 0.0 | 37 | 21.9 | 121 | 71.0 |
| Ashmore and Cartier Islands | 0 | 0.0 | 0 | 0.0 | 0 | 0.0 | 0 | 0.0 | 0 | 0.0 |
| Australia | 1496644 | 19.4 | 440791 | 5.7 | 2741193 | 35.5 | 3401622 | 44.0 | 618211 | 8.0 |
| Austria | 23984 | 28.6 | 14340 | 17.1 | 0 | 0.0 | 28141 | 33.5 | 0 | 0.0 |
| Azerbaijan | 8422 | 9.8 | 13583 | 15.7 | 0 | 0.0 | 16169 | 18.7 | 15574 | 18.0 |
| Bahamas | 4333 | 34.3 | 2890 | 22.8 | 1045 | 8.3 | 6196 | 49.0 | 5058 | 40.0 |
| Bahrain | 16 | 2.7 | 9 | 1.5 | 0 | 0.0 | 23 | 3.9 | 222 | 37.9 |
| Baikonur Cosmodrome | 0 | 0.0 | 0 | 0.0 | 0 | 0.0 | 0 | 0.0 | 0 | 0.0 |
| Bangladesh | 6221 | 4.5 | 6769 | 4.9 | 0 | 0.0 | 11614 | 8.4 | 22954 | 16.7 |
| Barbados | 9 | 1.9 | 66 | 14.8 | 0 | 0.0 | 74 | 16.5 | 323 | 72.3 |
| Belarus | 19392 | 9.4 | 16096 | 7.8 | 0 | 0.0 | 24254 | 11.7 | 11196 | 5.4 |
| Belgium | 7611 | 24.8 | 6136 | 20.0 | 0 | 0.0 | 9608 | 31.4 | 0 | 0.0 |
| Belize | 8377 | 37.4 | 14575 | 65.0 | 0 | 0.0 | 14763 | 65.8 | 1785 | 8.0 |
| Benin | 32848 | 28.1 | 15212 | 13.0 | 0 | 0.0 | 33716 | 28.9 | 8889 | 7.6 |
| Bermuda | 1 | 2.0 | 0 | 0.6 | 0 | 0.0 | 1 | 2.4 | 57 | 94.7 |
| Bhutan | 18847 | 46.5 | 13807 | 34.1 | 0 | 0.0 | 21348 | 52.7 | 4672 | 11.5 |
| Bolivia | 330881 | 30.3 | 233831 | 21.4 | 109865 | 10.1 | 475604 | 43.5 | 91069 | 8.3 |
| Bosnia and Herzegovina | 1252 | 2.4 | 1580 | 3.0 | 0 | 0.0 | 1976 | 3.8 | 7228 | 13.9 |
| Botswana | 170622 | 29.3 | 141530 | 24.3 | 113031 | 19.4 | 249242 | 42.8 | 3597 | 0.6 |
| Brazil | 2578317 | 30.2 | 962826 | 11.3 | 2013630 | 23.6 | 3556072 | 41.7 | 649944 | 7.6 |
| British Indian Ocean Territory | 49 | 100.0 | 1 | 2.7 | 0 | 0.0 | 49 | 100.0 | 0 | 0.0 |
| British Virgin Islands | 13 | 8.7 | 50 | 34.4 | 0 | 0.0 | 51 | 34.8 | 82 | 56.4 |
| Brunei Darussalam | 2446 | 42.5 | 1626 | 28.3 | 0 | 0.0 | 2892 | 50.3 | 893 | 15.5 |
| Bulgaria | 45949 | 40.7 | 26495 | 23.5 | 0 | 0.0 | 46724 | 41.4 | 0 | 0.0 |
| Burkina Faso | 42062 | 15.3 | 12854 | 4.7 | 0 | 0.0 | 44055 | 16.1 | 0 | 0.0 |
| Burundi | 1981 | 7.3 | 1611 | 5.9 | 0 | 0.0 | 2409 | 8.8 | 6347 | 23.3 |
| Cambodia | 47080 | 25.8 | 55672 | 30.6 | 6176 | 3.4 | 74362 | 40.8 | 17723 | 9.7 |
| Cameroon | 53198 | 11.4 | 40515 | 8.7 | 0 | 0.0 | 65019 | 13.9 | 100944 | 21.6 |
| Canada | 1090115 | 11.0 | 249719 | 2.5 | 8055412 | 81.2 | 8221501 | 82.9 | 85242 | 0.9 |
| Cape Verde | 117 | 3.0 | 629 | 16.1 | 0 | 0.0 | 640 | 16.4 | 3250 | 83.2 |
| Cayman Islands | 50 | 16.2 | 61 | 19.7 | 0 | 0.0 | 94 | 30.2 | 207 | 66.5 |
| Central African Republic | 111326 | 17.9 | 76691 | 12.3 | 139272 | 22.4 | 217621 | 35.0 | 31981 | 5.1 |
| Chad | 251777 | 19.8 | 127391 | 10.0 | 453372 | 35.6 | 684985 | 53.8 | 17976 | 1.4 |
| Chile | 143335 | 19.4 | 60257 | 8.2 | 148376 | 20.1 | 257395 | 34.9 | 170052 | 23.0 |
| China | 1600448 | 17.0 | 1125138 | 12.0 | 1192766 | 12.7 | 2595010 | 27.6 | 932678 | 9.9 |
| Clipperton Island | 0 | 0.0 | 5 | 100.0 | 0 | 0.0 | 5 | 100.0 | 0 | 0.0 |
| Colombia | 170033 | 14.9 | 100201 | 8.8 | 292352 | 25.6 | 431705 | 37.8 | 291093 | 25.5 |
| Comoros | 172 | 10.3 | 574 | 34.1 | 0 | 0.0 | 634 | 37.7 | 1039 | 61.8 |
| Cook Islands | 55 | 28.6 | 75 | 38.8 | 0 | 0.0 | 118 | 61.6 | 69 | 35.8 |
| Coral Sea Islands | 0 | 99.3 | 0 | 0.0 | 0 | 0.0 | 0 | 99.3 | 0 | 0.0 |
| Costa Rica | 14330 | 27.8 | 28415 | 55.2 | 0 | 0.0 | 29051 | 56.4 | 14956 | 29.1 |
| Cote d'Ivoire | 74390 | 23.0 | 23684 | 7.3 | 0 | 0.0 | 75860 | 23.5 | 15888 | 4.9 |
| Croatia | 20565 | 37.3 | 18116 | 32.9 | 0 | 0.0 | 23044 | 41.8 | 2234 | 4.1 |
| Cuba | 17087 | 15.5 | 20082 | 18.2 | 0 | 0.0 | 23431 | 21.2 | 50054 | 45.3 |
| Curacao | 65 | 13.9 | 105 | 22.5 | 0 | 0.0 | 125 | 26.7 | 281 | 60.2 |
| Cyprus | 3285 | 60.7 | 1843 | 34.1 | 0 | 0.0 | 3361 | 62.1 | 425 | 7.9 |
| Cyprus U.N. Buffer Zone | 49 | 14.2 | 29 | 8.5 | 0 | 0.0 | 55 | 16.2 | 80 | 23.3 |
| Czech Republic | 17371 | 22.1 | 7999 | 10.2 | 0 | 0.0 | 17502 | 22.2 | 0 | 0.0 |
| Dem. Rep. Korea | 3191 | 2.6 | 2668 | 2.2 | 0 | 0.0 | 5075 | 4.1 | 19895 | 16.2 |
| Democratic Republic of the Congo | 323661 | 13.8 | 162425 | 6.9 | 1174 | 0.1 | 373107 | 15.9 | 257888 | 11.0 |
| Denmark | 6513 | 15.3 | 3777 | 8.9 | 0 | 0.0 | 7560 | 17.8 | 2501 | 5.9 |
| Dhekelia | 14 | 11.1 | 1 | 0.9 | 0 | 0.0 | 15 | 11.5 | 11 | 8.8 |
| Djibouti | 223 | 1.0 | 1054 | 4.8 | 0 | 0.0 | 1256 | 5.7 | 6398 | 29.1 |
| Dominica | 170 | 23.1 | 218 | 29.7 | 0 | 0.0 | 287 | 39.1 | 446 | 60.7 |
| Dominican Republic | 12754 | 26.2 | 7551 | 15.5 | 0 | 0.0 | 13102 | 26.9 | 28222 | 58.0 |
| Ecuador | 55293 | 21.5 | 93773 | 36.5 | 30272 | 11.8 | 108757 | 42.4 | 100309 | 39.1 |
| Egypt | 128992 | 12.8 | 32720 | 3.3 | 473140 | 47.1 | 584014 | 58.1 | 31455 | 3.1 |
| El Salvador | 1733 | 8.4 | 2887 | 14.0 | 0 | 0.0 | 3361 | 16.3 | 9110 | 44.1 |
| Equatorial Guinea | 5067 | 18.9 | 3497 | 13.0 | 0 | 0.0 | 5069 | 18.9 | 2523 | 9.4 |
| Eritrea | 5917 | 4.8 | 14733 | 12.0 | 0 | 0.0 | 16592 | 13.5 | 13763 | 11.2 |
| Estonia | 9073 | 19.9 | 6488 | 14.2 | 0 | 0.0 | 9668 | 21.2 | 0 | 0.0 |
| Ethiopia | 199830 | 17.6 | 145802 | 12.9 | 4096 | 0.4 | 293336 | 25.9 | 129832 | 11.4 |
| Faeroe Islands | 8 | 0.6 | 9 | 0.7 | 0 | 0.0 | 16 | 1.2 | 1072 | 81.0 |
| Falkland Islands | 23 | 0.2 | 436 | 3.8 | 3210 | 27.7 | 3654 | 31.6 | 3522 | 30.4 |
| Federated States of Micronesia | 0 | 0.0 | 228 | 35.7 | 0 | 0.0 | 228 | 35.7 | 373 | 58.5 |
| Fiji | 1194 | 6.3 | 5047 | 26.5 | 0 | 0.0 | 5648 | 29.7 | 10598 | 55.7 |
| Finland | 43668 | 13.2 | 19896 | 6.0 | 45775 | 13.8 | 65458 | 19.7 | 13556 | 4.1 |
| France | 192055 | 30.1 | 72557 | 11.4 | 49769 | 7.8 | 217470 | 34.1 | 12090 | 1.9 |
| French Polynesia | 72 | 2.2 | 2584 | 77.0 | 0 | 0.0 | 2598 | 77.4 | 742 | 22.1 |
| French Southern and Antarctic Lands | 7269 | 99.9 | 1455 | 20.0 | 0 | 0.0 | 7269 | 99.9 | 0 | 0.0 |
| Gabon | 59511 | 22.7 | 27119 | 10.4 | 3900 | 1.5 | 64925 | 24.8 | 20215 | 7.7 |
| Georgia | 6329 | 9.1 | 20174 | 29.0 | 0 | 0.0 | 20985 | 30.1 | 14670 | 21.1 |
| Germany | 133175 | 37.3 | 56917 | 15.9 | 0 | 0.0 | 149362 | 41.8 | 526 | 0.1 |
| Ghana | 36182 | 15.1 | 18868 | 7.9 | 0 | 0.0 | 38730 | 16.1 | 40173 | 16.7 |
| Gibraltar | 2 | 61.0 | 4 | 97.5 | 0 | 0.0 | 4 | 97.9 | 0 | 0.0 |
| Greece | 45739 | 34.8 | 30839 | 23.4 | 0 | 0.0 | 51018 | 38.8 | 13522 | 10.3 |
| Greenland | 884783 | 41.3 | 15753 | 0.7 | 2074669 | 96.8 | 2099747 | 98.0 | 11633 | 0.5 |
| Grenada | 33 | 9.5 | 33 | 9.5 | 0 | 0.0 | 46 | 13.2 | 300 | 85.9 |
| Guam | 128 | 22.5 | 44 | 7.7 | 0 | 0.0 | 131 | 23.1 | 436 | 76.9 |
| Guatemala | 21838 | 20.0 | 53223 | 48.6 | 0 | 0.0 | 56021 | 51.2 | 30849 | 28.2 |
| Guernsey | 1 | 0.9 | 7 | 10.1 | 0 | 0.0 | 8 | 10.7 | 0 | 0.0 |
| Guinea | 85329 | 34.7 | 6518 | 2.7 | 0 | 0.0 | 87118 | 35.4 | 34888 | 14.2 |
| Guinea-Bissau | 5207 | 15.8 | 6789 | 20.6 | 0 | 0.0 | 9224 | 27.9 | 5762 | 17.4 |
| Guyana | 18553 | 8.7 | 1350 | 0.6 | 103911 | 48.9 | 107136 | 50.4 | 6260 | 2.9 |
| Haiti | 1924 | 7.1 | 5083 | 18.8 | 0 | 0.0 | 6057 | 22.4 | 19255 | 71.2 |
| Heard I. and McDonald Islands | 394 | 100.0 | 367 | 93.0 | 0 | 0.0 | 394 | 100.0 | 0 | 0.0 |
| Honduras | 26959 | 23.9 | 12455 | 11.0 | 3802 | 3.4 | 29486 | 26.1 | 44172 | 39.1 |
| Hong Kong | 464 | 44.5 | 683 | 65.6 | 0 | 0.0 | 718 | 68.9 | 184 | 17.7 |
| Hungary | 21088 | 22.6 | 14099 | 15.1 | 0 | 0.0 | 23422 | 25.1 | 0 | 0.0 |
| Iceland | 19880 | 19.5 | 9636 | 9.5 | 38633 | 37.9 | 49442 | 48.5 | 4381 | 4.3 |
| India | 183890 | 5.8 | 213088 | 6.7 | 17170 | 0.5 | 326555 | 10.3 | 462776 | 14.6 |
| Indian Ocean Territories | 40 | 35.7 | 65 | 57.6 | 0 | 0.0 | 66 | 58.2 | 47 | 41.8 |
| Indonesia | 230337 | 12.2 | 282336 | 14.9 | 151865 | 8.0 | 496929 | 26.3 | 418932 | 22.1 |
| Iran | 139209 | 8.6 | 100859 | 6.2 | 8385 | 0.5 | 173364 | 10.7 | 204979 | 12.6 |
| Iraq | 6680 | 1.5 | 30587 | 7.0 | 1386 | 0.3 | 31111 | 7.1 | 37889 | 8.6 |
| Ireland | 9778 | 14.1 | 3445 | 5.0 | 0 | 0.0 | 11192 | 16.1 | 1056 | 1.5 |
| Isle of Man | 37 | 6.5 | 176 | 30.7 | 0 | 0.0 | 197 | 34.3 | 0 | 0.0 |
| Israel | 4998 | 22.8 | 7926 | 36.1 | 0 | 0.0 | 11194 | 51.0 | 6327 | 28.8 |
| Italy | 64732 | 21.5 | 60059 | 19.9 | 0 | 0.0 | 79980 | 26.5 | 43651 | 14.5 |
| Jamaica | 1800 | 16.2 | 2915 | 26.3 | 0 | 0.0 | 3188 | 28.7 | 7629 | 68.8 |
| Japan | 109969 | 29.4 | 61281 | 16.4 | 0 | 0.0 | 131883 | 35.2 | 30535 | 8.2 |
| Jersey | 17 | 14.5 | 5 | 3.9 | 0 | 0.0 | 19 | 16.4 | 0 | 0.0 |
| Jordan | 2828 | 3.2 | 8166 | 9.2 | 0 | 0.0 | 8912 | 10.0 | 10722 | 12.0 |
| Kazakhstan | 90662 | 3.3 | 153786 | 5.7 | 11987 | 0.4 | 229310 | 8.5 | 218069 | 8.0 |
| Kenya | 73657 | 12.5 | 70492 | 12.0 | 9099 | 1.5 | 111319 | 18.9 | 85379 | 14.5 |
| Kiribati | 141 | 14.8 | 652 | 68.2 | 0 | 0.0 | 652 | 68.2 | 197 | 20.6 |
| Kosovo | 65 | 0.6 | 2596 | 23.8 | 0 | 0.0 | 2605 | 23.9 | 885 | 8.1 |
| Kuwait | 2797 | 15.9 | 1025 | 5.8 | 0 | 0.0 | 2894 | 16.5 | 1256 | 7.2 |
| Kyrgyzstan | 14326 | 7.2 | 6231 | 3.1 | 68 | 0.0 | 19037 | 9.6 | 34046 | 17.1 |
| Lao PDR | 38121 | 16.6 | 50761 | 22.1 | 7 | 0.0 | 56813 | 24.8 | 57631 | 25.1 |
| Latvia | 11683 | 18.1 | 5380 | 8.4 | 0 | 0.0 | 11762 | 18.3 | 0 | 0.0 |
| Lebanon | 274 | 2.7 | 3379 | 33.7 | 0 | 0.0 | 3456 | 34.5 | 3560 | 35.5 |
| Lesotho | 81 | 0.3 | 2440 | 8.1 | 0 | 0.0 | 2450 | 8.1 | 10002 | 33.1 |
| Liberia | 4010 | 4.2 | 27852 | 29.0 | 0 | 0.0 | 28946 | 30.2 | 16610 | 17.3 |
| Libya | 4208 | 0.3 | 37162 | 2.3 | 946245 | 58.0 | 985103 | 60.4 | 9216 | 0.6 |
| Liechtenstein | 48 | 34.7 | 4 | 3.1 | 0 | 0.0 | 48 | 34.9 | 0 | 0.0 |
| Lithuania | 11162 | 17.2 | 5675 | 8.8 | 0 | 0.0 | 11518 | 17.8 | 889 | 1.4 |
| Luxembourg | 1297 | 49.8 | 451 | 17.3 | 0 | 0.0 | 1349 | 51.8 | 0 | 0.0 |
| Macao | 0 | 0.0 | 20 | 66.0 | 0 | 0.0 | 20 | 66.0 | 1 | 3.4 |
| Macedonia | 2865 | 11.3 | 8476 | 33.4 | 0 | 0.0 | 9325 | 36.7 | 1691 | 6.7 |
| Madagascar | 33242 | 5.6 | 96063 | 16.1 | 0 | 0.0 | 107094 | 18.0 | 208259 | 34.9 |
| Malawi | 27006 | 22.5 | 19630 | 16.3 | 612 | 0.5 | 31106 | 25.9 | 25770 | 21.5 |
| Malaysia | 60777 | 18.4 | 54799 | 16.6 | 6687 | 2.0 | 95431 | 28.9 | 110549 | 33.5 |
| Maldives | 1 | 1.2 | 1 | 0.9 | 0 | 0.0 | 2 | 2.0 | 105 | 95.7 |
| Mali | 104708 | 8.3 | 24630 | 2.0 | 523805 | 41.6 | 633574 | 50.3 | 31693 | 2.5 |
| Malta | 107 | 32.9 | 42 | 12.7 | 0 | 0.0 | 109 | 33.5 | 161 | 49.4 |
| Marshall Islands | 20 | 11.9 | 12 | 7.0 | 0 | 0.0 | 30 | 18.3 | 68 | 41.0 |
| Mauritania | 6247 | 0.6 | 10740 | 1.0 | 646704 | 62.1 | 655421 | 62.9 | 28112 | 2.7 |
| Mauritius | 107 | 5.3 | 608 | 30.0 | 0 | 0.0 | 644 | 31.8 | 1294 | 63.9 |
| Mexico | 285671 | 14.5 | 328742 | 16.7 | 5180 | 0.3 | 523708 | 26.6 | 464896 | 23.6 |
| Moldova | 1268 | 3.8 | 879 | 2.6 | 0 | 0.0 | 1908 | 5.7 | 457 | 1.4 |
| Monaco | 5 | 29.1 | 0 | 0.0 | 0 | 0.0 | 5 | 29.1 | 9 | 49.5 |
| Mongolia | 275793 | 17.6 | 82285 | 5.3 | 265644 | 17.0 | 446574 | 28.6 | 31403 | 2.0 |
| Montenegro | 1235 | 9.0 | 1526 | 11.1 | 0 | 0.0 | 2252 | 16.4 | 4832 | 35.2 |
| Montserrat | 11 | 11.2 | 16 | 15.7 | 0 | 0.0 | 17 | 17.3 | 83 | 82.7 |
| Morocco | 137847 | 23.2 | 56098 | 9.4 | 29721 | 5.0 | 175337 | 29.5 | 46400 | 7.8 |
| Mozambique | 170089 | 21.5 | 32848 | 4.1 | 12983 | 1.6 | 189613 | 23.9 | 63856 | 8.1 |
| Myanmar | 42301 | 6.3 | 114668 | 17.2 | 6726 | 1.0 | 120505 | 18.1 | 127648 | 19.2 |
| Namibia | 312298 | 37.8 | 105249 | 12.7 | 144681 | 17.5 | 365022 | 44.2 | 58917 | 7.1 |
| Nauru | 0 | 0.0 | 1 | 2.1 | 0 | 0.0 | 1 | 2.1 | 28 | 97.9 |
| Nepal | 33555 | 22.7 | 34200 | 23.2 | 406 | 0.3 | 40640 | 27.5 | 32673 | 22.1 |
| Netherlands | 10001 | 26.8 | 5373 | 14.4 | 671 | 1.8 | 10350 | 27.7 | 3337 | 8.9 |
| New Caledonia | 11201 | 59.2 | 4109 | 21.7 | 0 | 0.0 | 12539 | 66.2 | 6144 | 32.4 |
| New Zealand | 86145 | 32.1 | 12726 | 4.7 | 18085 | 6.7 | 97824 | 36.4 | 57307 | 21.3 |
| Nicaragua | 48063 | 37.1 | 24021 | 18.6 | 0 | 0.0 | 49174 | 38.0 | 20331 | 15.7 |
| Niger | 209231 | 17.6 | 99833 | 8.4 | 626788 | 52.8 | 695032 | 58.5 | 0 | 0.0 |
| Nigeria | 126553 | 13.9 | 35960 | 3.9 | 0 | 0.0 | 129995 | 14.2 | 80833 | 8.9 |
| Niue | 39 | 17.5 | 31 | 13.9 | 0 | 0.0 | 41 | 18.4 | 0 | 0.0 |
| Norfolk Island | 12 | 28.8 | 41 | 100.0 | 0 | 0.0 | 41 | 100.0 | 0 | 0.0 |
| Northern Cyprus | 44 | 1.3 | 671 | 20.2 | 0 | 0.0 | 715 | 21.5 | 1089 | 32.7 |
| Northern Mariana Islands | 62 | 10.6 | 208 | 35.7 | 0 | 0.0 | 230 | 39.4 | 352 | 60.4 |
| Norway | 95950 | 25.2 | 55245 | 14.5 | 92247 | 24.3 | 148670 | 39.1 | 16615 | 4.4 |
| Oman | 8066 | 2.6 | 68930 | 22.0 | 100880 | 32.3 | 128443 | 41.1 | 29200 | 9.3 |
| Pakistan | 96219 | 11.0 | 55036 | 6.3 | 1507 | 0.2 | 118340 | 13.5 | 110954 | 12.7 |
| Palau | 224 | 46.5 | 316 | 65.7 | 0 | 0.0 | 429 | 89.1 | 52 | 10.7 |
| Palestine | 550 | 8.7 | 1797 | 28.5 | 0 | 0.0 | 1832 | 29.1 | 2800 | 44.5 |
| Panama | 15275 | 20.4 | 21300 | 28.4 | 0 | 0.0 | 22855 | 30.5 | 37127 | 49.5 |
| Papua New Guinea | 17738 | 3.8 | 70734 | 15.1 | 3 | 0.0 | 75403 | 16.1 | 190673 | 40.7 |
| Paraguay | 57965 | 14.4 | 35712 | 8.9 | 64599 | 16.1 | 115738 | 28.8 | 40083 | 10.0 |
| Peru | 277796 | 21.4 | 227084 | 17.5 | 272107 | 21.0 | 502780 | 38.7 | 298928 | 23.0 |
| Philippines | 45365 | 15.4 | 70980 | 24.1 | 0 | 0.0 | 87884 | 29.8 | 102886 | 34.9 |
| Pitcairn Islands | 36 | 85.0 | 36 | 83.4 | 0 | 0.0 | 42 | 99.3 | 0 | 0.3 |
| Poland | 124359 | 39.7 | 53609 | 17.1 | 0 | 0.0 | 129131 | 41.3 | 3319 | 1.1 |
| Portugal | 20429 | 22.3 | 13481 | 14.7 | 0 | 0.0 | 22580 | 24.7 | 4372 | 4.8 |
| Puerto Rico | 619 | 6.8 | 1783 | 19.7 | 0 | 0.0 | 1830 | 20.2 | 6235 | 68.9 |
| Qatar | 1158 | 10.3 | 47 | 0.4 | 0 | 0.0 | 1176 | 10.5 | 2840 | 25.4 |
| Republic of Congo | 133497 | 38.4 | 68866 | 19.8 | 8737 | 2.5 | 153819 | 44.3 | 24756 | 7.1 |
| Republic of Korea | 15917 | 16.1 | 464 | 0.5 | 0 | 0.0 | 16301 | 16.5 | 25447 | 25.8 |
| Romania | 56080 | 23.7 | 38121 | 16.1 | 0 | 0.0 | 63419 | 26.8 | 13746 | 5.8 |
| Russian Federation | 1641637 | 9.7 | 895221 | 5.3 | 9399338 | 55.6 | 10311203 | 61.0 | 175970 | 1.0 |
| Rwanda | 2382 | 9.3 | 5954 | 23.4 | 0 | 0.0 | 5992 | 23.5 | 7008 | 27.5 |
| Saint Helena | 95 | 25.9 | 127 | 34.5 | 0 | 0.0 | 201 | 54.6 | 156 | 42.4 |
| Saint Kitts and Nevis | 70 | 26.4 | 63 | 23.8 | 0 | 0.0 | 93 | 35.0 | 168 | 63.3 |
| Saint Lucia | 118 | 19.3 | 212 | 34.8 | 0 | 0.0 | 220 | 36.2 | 382 | 62.8 |
| Saint Pierre and Miquelon | 7 | 2.9 | 16 | 6.6 | 0 | 0.0 | 23 | 9.5 | 0 | 0.0 |
| Saint Vincent and the Grenadines | 84 | 22.6 | 140 | 37.7 | 0 | 0.0 | 146 | 39.4 | 221 | 59.7 |
| Saint-Bartholemy | 10 | 38.7 | 7 | 27.7 | 0 | 0.0 | 14 | 58.6 | 10 | 41.3 |
| Saint-Martin | 18 | 26.8 | 2 | 3.0 | 0 | 0.0 | 20 | 29.8 | 45 | 66.1 |
| Samoa | 222 | 7.9 | 1027 | 36.7 | 0 | 0.0 | 1155 | 41.3 | 1278 | 45.7 |
| San Marino | 0 | 0.0 | 0 | 0.0 | 0 | 0.0 | 0 | 0.0 | 0 | 0.0 |
| Sao Tome and Principe | 311 | 29.8 | 535 | 51.3 | 0 | 0.0 | 548 | 52.5 | 482 | 46.2 |
| Saudi Arabia | 92149 | 4.8 | 48260 | 2.5 | 0 | 0.0 | 109743 | 5.7 | 139927 | 7.2 |
| Scarborough Reef | 0 | 0.0 | 0 | 0.0 | 0 | 0.0 | 0 | 0.0 | 0 | 0.0 |
| Senegal | 50201 | 25.4 | 28463 | 14.4 | 0 | 0.0 | 55059 | 27.9 | 20497 | 10.4 |
| Serbia | 6892 | 8.9 | 10315 | 13.3 | 0 | 0.0 | 12240 | 15.8 | 6 | 0.0 |
| Serranilla Bank | 0 | 0.0 | 0 | 0.0 | 0 | 0.0 | 0 | 0.0 | 0 | 0.0 |
| Seychelles | 190 | 46.1 | 255 | 61.9 | 0 | 0.0 | 262 | 63.7 | 132 | 32.2 |
| Siachen Glacier | 11 | 0.5 | 0 | 0.0 | 937 | 44.8 | 938 | 44.8 | 0 | 0.0 |
| Sierra Leone | 7634 | 10.6 | 4510 | 6.3 | 0 | 0.0 | 9170 | 12.7 | 12950 | 18.0 |
| Singapore | 32 | 6.3 | 90 | 17.5 | 0 | 0.0 | 90 | 17.5 | 165 | 32.1 |
| Sint Maarten | 0 | 0.8 | 2 | 8.6 | 0 | 0.0 | 2 | 9.4 | 18 | 75.9 |
| Slovakia | 18012 | 37.2 | 13660 | 28.2 | 0 | 0.0 | 19204 | 39.7 | 0 | 0.0 |
| Slovenia | 10843 | 53.4 | 5585 | 27.5 | 0 | 0.0 | 11223 | 55.2 | 252 | 1.2 |
| Solomon Islands | 536 | 2.0 | 12711 | 46.4 | 0 | 0.0 | 12829 | 46.9 | 10110 | 36.9 |
| Somalia | 82 | 0.0 | 33821 | 7.1 | 0 | 0.0 | 33823 | 7.1 | 100206 | 21.1 |
| Somaliland | 123 | 0.1 | 15379 | 9.1 | 0 | 0.0 | 15500 | 9.2 | 36311 | 21.6 |
| South Africa | 101086 | 8.3 | 187649 | 15.3 | 940 | 0.1 | 224475 | 18.3 | 196392 | 16.0 |
| South Georgia and South Sandwich Islands | 535 | 13.7 | 3855 | 98.5 | 0 | 0.0 | 3913 | 100.0 | 0 | 0.0 |
| South Sudan | 103466 | 16.4 | 89665 | 14.2 | 136 | 0.0 | 130539 | 20.7 | 20188 | 3.2 |
| Spain | 143003 | 28.2 | 183681 | 36.2 | 0 | 0.0 | 225326 | 44.4 | 27357 | 5.4 |
| Spratly Islands | 0 | 0.0 | 0 | 0.0 | 0 | 0.0 | 0 | 0.0 | 0 | 0.0 |
| Sri Lanka | 19900 | 29.8 | 6068 | 9.1 | 0 | 0.0 | 22798 | 34.2 | 28269 | 42.4 |
| Sudan | 38155 | 2.0 | 55330 | 3.0 | 151716 | 8.1 | 234786 | 12.6 | 73461 | 3.9 |
| Suriname | 21598 | 14.8 | 46879 | 32.1 | 110485 | 75.6 | 118726 | 81.3 | 4448 | 3.0 |
| Swaziland | 754 | 4.4 | 2786 | 16.2 | 0 | 0.0 | 2923 | 17.0 | 7410 | 43.1 |
| Sweden | 62941 | 14.2 | 17532 | 3.9 | 50820 | 11.4 | 94212 | 21.2 | 9017 | 2.0 |
| Switzerland | 4345 | 10.5 | 5491 | 13.3 | 0 | 0.0 | 8699 | 21.0 | 261 | 0.6 |
| Syria | 1294 | 0.7 | 18654 | 10.0 | 0 | 0.0 | 19265 | 10.3 | 16005 | 8.6 |
| Taiwan | 7130 | 19.6 | 6639 | 18.3 | 0 | 0.0 | 8351 | 23.0 | 20994 | 57.7 |
| Tajikistan | 31165 | 21.9 | 13009 | 9.1 | 28088 | 19.7 | 52600 | 36.9 | 17357 | 12.2 |
| Tanzania | 363188 | 38.3 | 185685 | 19.6 | 1094 | 0.1 | 406454 | 42.9 | 115038 | 12.1 |
| Thailand | 96001 | 18.6 | 85458 | 16.5 | 0 | 0.0 | 121716 | 23.5 | 139611 | 27.0 |
| The Gambia | 444 | 4.2 | 741 | 7.0 | 0 | 0.0 | 816 | 7.7 | 7905 | 74.8 |
| Timor-Leste | 2363 | 15.6 | 3817 | 25.1 | 0 | 0.0 | 4367 | 28.8 | 4141 | 27.3 |
| Togo | 14224 | 24.9 | 4775 | 8.3 | 0 | 0.0 | 14361 | 25.1 | 4432 | 7.7 |
| Tonga | 62 | 10.3 | 166 | 27.3 | 0 | 0.0 | 189 | 31.1 | 243 | 40.1 |
| Trinidad and Tobago | 1594 | 30.9 | 988 | 19.2 | 0 | 0.0 | 1809 | 35.1 | 2728 | 52.9 |
| Tunisia | 12388 | 7.9 | 11269 | 7.2 | 2877 | 1.8 | 16083 | 10.2 | 11449 | 7.3 |
| Turkey | 1772 | 0.2 | 155771 | 19.9 | 3120 | 0.4 | 158080 | 20.2 | 93474 | 12.0 |
| Turkmenistan | 15194 | 3.2 | 32172 | 6.8 | 785 | 0.2 | 40932 | 8.7 | 49084 | 10.4 |
| Turks and Caicos Islands | 109 | 24.3 | 152 | 34.0 | 0 | 0.0 | 204 | 45.6 | 241 | 53.8 |
| Tuvalu | 2 | 10.5 | 0 | 0.0 | 0 | 0.0 | 2 | 10.5 | 0 | 0.0 |
| Uganda | 38835 | 15.9 | 23976 | 9.8 | 2212 | 0.9 | 44672 | 18.3 | 42125 | 17.3 |
| Ukraine | 24842 | 4.2 | 26839 | 4.5 | 0 | 0.0 | 46446 | 7.8 | 51475 | 8.6 |
| United Arab Emirates | 12674 | 17.7 | 14 | 0.0 | 0 | 0.0 | 12688 | 17.8 | 11842 | 16.6 |
| United Kingdom | 69357 | 28.5 | 23782 | 9.8 | 0 | 0.0 | 71565 | 29.4 | 11247 | 4.6 |
| United States | 1223937 | 12.9 | 398063 | 4.2 | 1606138 | 17.0 | 2398673 | 25.3 | 1163547 | 12.3 |
| United States Minor Outlying Islands | 25 | 100.0 | 25 | 100.0 | 0 | 0.0 | 25 | 100.0 | 0 | 0.0 |
| United States Virgin Islands | 59 | 16.5 | 84 | 23.5 | 0 | 0.0 | 99 | 27.8 | 199 | 55.9 |
| Uruguay | 6128 | 3.4 | 34256 | 19.3 | 303 | 0.2 | 35042 | 19.7 | 16561 | 9.3 |
| US Naval Base Guantanamo Bay | 0 | 0.0 | 0 | 0.0 | 0 | 0.0 | 0 | 0.0 | 66 | 100.0 |
| Uzbekistan | 14523 | 3.2 | 32738 | 7.3 | 24607 | 5.5 | 64332 | 14.4 | 42431 | 9.5 |
| Vanuatu | 513 | 4.1 | 4175 | 33.8 | 0 | 0.0 | 4403 | 35.6 | 4753 | 38.4 |
| Venezuela | 490050 | 53.3 | 224384 | 24.4 | 303341 | 33.0 | 581639 | 63.3 | 69972 | 7.6 |
| Vietnam | 26560 | 8.0 | 38099 | 11.5 | 199 | 0.1 | 47777 | 14.4 | 158404 | 47.9 |
| Wallis and Futuna Islands | 0 | 0.0 | 18 | 12.8 | 0 | 0.0 | 18 | 12.8 | 6 | 4.1 |
| Western Sahara | 218 | 0.2 | 218 | 0.2 | 23954 | 26.4 | 24161 | 26.6 | 20 | 0.0 |
| Yemen | 3479 | 0.8 | 17285 | 3.8 | 77267 | 17.0 | 96967 | 21.3 | 60631 | 13.3 |
| Zambia | 311917 | 41.2 | 104547 | 13.8 | 2613 | 0.3 | 331082 | 43.8 | 19506 | 2.6 |
| Zimbabwe | 105686 | 27.0 | 31923 | 8.2 | 0 | 0.0 | 107297 | 27.4 | 12541 | 3.2 |

| ***Extended Data Table 3.*** *The area and proportion of intact land requiring conservation within each country that is at risk of habitat conversion by 2030 and 2050 based on Shared Socioeconomic Pathways 1 (optimistic) and 2 (middle of the road).* | | | | | | | | |
| --- | --- | --- | --- | --- | --- | --- | --- | --- |
| **Country** | **Habitat loss by 2030 (SSP1) (km2)** | **Habitat loss by 2030 (SSP1) (%)** | **Habitat loss by 2050 (SSP1) (km2)** | **Habitat loss by 2050 (SSP1) (%)** | **Habitat loss by 2030 (SSP2) (km2)** | **Habitat loss by 2030 (SSP2) (%)** | **Habitat loss by 2050 (SSP2) (km2)** | **Habitat loss by 2050 (SSP2) (%)** |
| Afghanistan | 155 | 0.27 | 136 | 0.23 | 194 | 0.34 | 356 | 0.62 |
| Akrotiri | 0 | 0.00 | 0 | 0.00 | 0 | 0.00 | 0 | 0.00 |
| Aland Islands | 0 | 0.00 | 0 | 0.00 | 0 | 0.00 | 0 | 0.00 |
| Albania | 16 | 0.31 | 66 | 1.26 | 55 | 1.05 | 168 | 3.21 |
| Algeria | 708 | 0.06 | 3895 | 0.32 | 6772 | 0.56 | 12467 | 1.03 |
| American Samoa | 0 | 0.00 | 0 | 0.00 | 0 | 0.00 | 0 | 0.00 |
| Andorra | 0 | 0.00 | 0 | 0.00 | 5 | 0.69 | 14 | 2.14 |
| Angola | 1051 | 0.75 | 1340 | 0.96 | 3855 | 2.75 | 8454 | 6.04 |
| Anguilla | 0 | 0.00 | 0 | 0.00 | 0 | 0.00 | 0 | 0.00 |
| Antigua and Barbuda | 0 | 0.00 | 0 | 0.00 | 0 | 0.00 | 0 | 0.00 |
| Argentina | 2515 | 0.50 | 6373 | 1.28 | 24724 | 4.95 | 45400 | 9.09 |
| Armenia | 15 | 0.23 | 100 | 1.53 | 647 | 9.85 | 1418 | 21.59 |
| Aruba | 0 | 0.00 | 0 | 0.00 | 0 | 0.00 | 0 | 0.00 |
| Ashmore and Cartier Islands | 0 | 0.00 | 0 | 0.00 | 0 | 0.00 | 0 | 0.00 |
| Australia | 2135 | 0.08 | 26946 | 1.06 | 42392 | 1.66 | 72219 | 2.83 |
| Austria | 17 | 0.10 | 45 | 0.27 | 194 | 1.15 | 204 | 1.21 |
| Azerbaijan | 103 | 0.70 | 491 | 3.32 | 1774 | 12.01 | 3604 | 24.39 |
| Bahamas | 7 | 0.17 | 6 | 0.15 | 358 | 8.92 | 549 | 13.69 |
| Bahrain | 7 | 1.66 | 20 | 4.55 | 18 | 4.00 | 46 | 10.32 |
| Baikonur Cosmodrome | 0 | 0.00 | 0 | 0.00 | 0 | 0.00 | 0 | 0.00 |
| Bangladesh | 265 | 1.77 | 365 | 2.44 | 556 | 3.71 | 885 | 5.90 |
| Barbados | 14 | 7.12 | 9 | 4.47 | 13 | 6.26 | 31 | 15.45 |
| Belarus | 9 | 0.04 | 165 | 0.82 | 573 | 2.85 | 886 | 4.41 |
| Belgium | 12 | 0.21 | 16 | 0.27 | 323 | 5.54 | 733 | 12.58 |
| Belize | 11 | 0.06 | 114 | 0.64 | 624 | 3.49 | 754 | 4.22 |
| Benin | 2656 | 9.93 | 3531 | 13.20 | 2425 | 9.06 | 7234 | 27.04 |
| Bermuda | 0 | 0.00 | 0 | 0.00 | 0 | 0.00 | 0 | 0.00 |
| Bhutan | 120 | 0.59 | 82 | 0.40 | 331 | 1.62 | 654 | 3.20 |
| Bolivia | 1676 | 0.42 | 2211 | 0.56 | 17389 | 4.39 | 28757 | 7.26 |
| Bosnia and Herzegovina | 12 | 0.29 | 68 | 1.72 | 146 | 3.67 | 153 | 3.85 |
| Botswana | 49 | 0.03 | 107 | 0.07 | 2704 | 1.78 | 4674 | 3.08 |
| Brazil | 15392 | 0.45 | 40833 | 1.19 | 108395 | 3.15 | 169138 | 4.92 |
| British Indian Ocean Territory | 0 | 0.00 | 0 | 0.00 | 0 | 0.00 | 0 | 0.00 |
| British Virgin Islands | 0 | 0.00 | 0 | 0.00 | 0 | 0.00 | 0 | 0.00 |
| Brunei Darussalam | 15 | 0.31 | 52 | 1.08 | 39 | 0.80 | 50 | 1.03 |
| Bulgaria | 14 | 0.06 | 93 | 0.37 | 1076 | 4.26 | 1645 | 6.51 |
| Burkina Faso | 2780 | 8.91 | 2375 | 7.61 | 1926 | 6.17 | 3255 | 10.43 |
| Burundi | 39 | 2.27 | 86 | 4.92 | 110 | 6.33 | 112 | 6.44 |
| Cambodia | 386 | 0.57 | 1411 | 2.07 | 3824 | 5.61 | 7259 | 10.65 |
| Cameroon | 2745 | 2.16 | 4476 | 3.53 | 5024 | 3.96 | 11323 | 8.92 |
| Canada | 552 | 0.01 | 1112 | 0.02 | 25262 | 0.36 | 44646 | 0.63 |
| Cape Verde | 17 | 1.23 | 0 | 0.00 | 125 | 8.94 | 273 | 19.53 |
| Cayman Islands | 0 | 0.00 | 0 | 0.00 | 0 | 0.00 | 0 | 0.00 |
| Central African Republic | 1438 | 0.62 | 2180 | 0.94 | 2540 | 1.10 | 5536 | 2.40 |
| Chad | 1621 | 0.30 | 1605 | 0.30 | 15931 | 2.99 | 33932 | 6.37 |
| Chile | 171 | 0.06 | 199 | 0.07 | 9077 | 3.14 | 12401 | 4.29 |
| China | 5644 | 0.37 | 6228 | 0.41 | 34815 | 2.27 | 60955 | 3.97 |
| Clipperton Island | 0 | 0.00 | 0 | 0.00 | 0 | 0.00 | 0 | 0.00 |
| Colombia | 3053 | 0.59 | 4522 | 0.87 | 13410 | 2.57 | 21475 | 4.12 |
| Comoros | 0 | 0.00 | 0 | 0.00 | 0 | 87.13 | 0 | 100.00 |
| Cook Islands | 0 | 0.00 | 0 | 0.00 | 0 | 0.00 | 0 | 0.00 |
| Coral Sea Islands | 0 | 0.00 | 0 | 0.00 | 0 | 0.00 | 0 | 0.00 |
| Costa Rica | 53 | 0.24 | 104 | 0.47 | 894 | 4.05 | 1476 | 6.69 |
| Cote d'Ivoire | 831 | 2.13 | 1354 | 3.46 | 2146 | 5.49 | 4314 | 11.03 |
| Croatia | 76 | 0.44 | 217 | 1.24 | 708 | 4.07 | 1095 | 6.29 |
| Cuba | 10 | 0.04 | 12 | 0.05 | 3502 | 14.44 | 5480 | 22.60 |
| Curacao | 0 | 0.00 | 0 | 0.00 | 0 | 0.00 | 0 | 0.00 |
| Cyprus | 11 | 0.26 | 18 | 0.43 | 104 | 2.46 | 199 | 4.69 |
| Cyprus U.N. Buffer Zone | 0 | 0.00 | 0 | 0.00 | 0 | 0.00 | 0 | 0.00 |
| Czech Republic | 0 | 0.00 | 0 | 0.00 | 150 | 1.99 | 298 | 3.96 |
| Dem. Rep. Korea | 33 | 0.21 | 11 | 0.07 | 558 | 3.50 | 773 | 4.85 |
| Democratic Republic of the Congo | 7930 | 1.44 | 10074 | 1.83 | 9350 | 1.70 | 17735 | 3.22 |
| Denmark | 2 | 0.08 | 4 | 0.18 | 152 | 6.06 | 132 | 5.29 |
| Dhekelia | 0 | 0.00 | 0 | 0.00 | 0 | 0.00 | 0 | 0.00 |
| Djibouti | 1 | 0.09 | 4 | 0.32 | 94 | 6.77 | 212 | 15.18 |
| Dominica | 0 | 0.00 | 0 | 0.00 | 0 | 0.00 | 0 | 0.00 |
| Dominican Republic | 9 | 0.05 | 254 | 1.58 | 2852 | 17.76 | 4151 | 25.84 |
| Ecuador | 995 | 0.68 | 1777 | 1.21 | 4974 | 3.38 | 9077 | 6.16 |
| Egypt | 1294 | 0.22 | 10113 | 1.71 | 6462 | 1.09 | 13752 | 2.33 |
| El Salvador | 14 | 0.65 | 98 | 4.40 | 404 | 18.15 | 589 | 26.45 |
| Equatorial Guinea | 31 | 0.52 | 49 | 0.83 | 27 | 0.46 | 30 | 0.50 |
| Eritrea | 101 | 0.98 | 295 | 2.86 | 2133 | 20.71 | 3965 | 38.50 |
| Estonia | 0 | 0.00 | 0 | 0.00 | 189 | 3.23 | 376 | 6.42 |
| Ethiopia | 5112 | 1.81 | 7436 | 2.64 | 21414 | 7.59 | 45837 | 16.25 |
| Faeroe Islands | 0 | 100.00 | 0 | 100.00 | 0 | 65.24 | 0 | 65.24 |
| Falkland Islands | 0 | 0.00 | 0 | 0.00 | 3 | 0.05 | 1 | 0.02 |
| Federated States of Micronesia | 0 | 0.00 | 0 | 0.00 | 0 | 0.00 | 0 | 0.00 |
| Fiji | 0 | 0.00 | 0 | 0.00 | 0 | 0.00 | 0 | 0.00 |
| Finland | 33 | 0.05 | 119 | 0.18 | 555 | 0.85 | 642 | 0.98 |
| France | 355 | 0.42 | 523 | 0.62 | 1996 | 2.36 | 4320 | 5.10 |
| French Polynesia | 0 | 0.00 | 0 | 0.00 | 0 | 0.00 | 0 | 0.00 |
| French Southern and Antarctic Lands | 0 | 0.00 | 0 | 0.00 | 0 | 0.00 | 0 | 0.00 |
| Gabon | 79 | 0.11 | 214 | 0.30 | 699 | 0.98 | 455 | 0.64 |
| Georgia | 22 | 0.10 | 278 | 1.27 | 1029 | 4.72 | 2949 | 13.53 |
| Germany | 80 | 0.10 | 101 | 0.13 | 3665 | 4.68 | 7912 | 10.09 |
| Ghana | 1212 | 5.32 | 1194 | 5.24 | 2349 | 10.31 | 4660 | 20.46 |
| Gibraltar | 0 | 0.00 | 0 | 0.00 | 0 | 0.00 | 0 | 0.00 |
| Greece | 15 | 0.08 | 85 | 0.45 | 989 | 5.19 | 2098 | 11.01 |
| Greenland | 1 | 0.00 | 2 | 0.00 | 273 | 0.12 | 856 | 0.37 |
| Grenada | 0 | 0.00 | 0 | 0.00 | 0 | 0.00 | 0 | 0.00 |
| Guam | 0 | 0.00 | 0 | 0.00 | 0 | 0.00 | 0 | 0.00 |
| Guatemala | 286 | 0.56 | 1811 | 3.54 | 3701 | 7.23 | 6779 | 13.25 |
| Guernsey | 0 | 0.00 | 0 | 0.00 | 0 | 0.00 | 0 | 0.00 |
| Guinea | 3845 | 8.04 | 4006 | 8.38 | 7949 | 16.63 | 16791 | 35.12 |
| Guinea-Bissau | 182 | 3.05 | 354 | 5.92 | 544 | 9.11 | 1038 | 17.39 |
| Guyana | 125 | 0.11 | 542 | 0.50 | 1989 | 1.83 | 2266 | 2.08 |
| Haiti | 0 | 0.00 | 0 | 0.00 | 825 | 12.93 | 762 | 11.94 |
| Heard I. and McDonald Islands | 0 | 0.00 | 0 | 0.00 | 0 | 0.00 | 0 | 0.00 |
| Honduras | 237 | 0.46 | 1126 | 2.18 | 4547 | 8.82 | 7452 | 14.46 |
| Hong Kong | 0 | 0.00 | 0 | 0.00 | 0 | 0.00 | 0 | 0.00 |
| Hungary | 27 | 0.31 | 100 | 1.13 | 220 | 2.50 | 260 | 2.95 |
| Iceland | 5 | 0.01 | 12 | 0.03 | 82 | 0.23 | 126 | 0.35 |
| India | 2333 | 0.55 | 5400 | 1.28 | 25350 | 6.01 | 42332 | 10.03 |
| Indian Ocean Territories | 0 | 0.00 | 0 | 0.00 | 0 | 0.00 | 0 | 0.00 |
| Indonesia | 1173 | 0.19 | 3270 | 0.53 | 16859 | 2.71 | 31485 | 5.06 |
| Iran | 3689 | 1.53 | 8061 | 3.35 | 12253 | 5.09 | 20761 | 8.63 |
| Iraq | 2006 | 3.64 | 3482 | 6.32 | 1586 | 2.88 | 3117 | 5.66 |
| Ireland | 58 | 1.03 | 75 | 1.33 | 101 | 1.79 | 62 | 1.11 |
| Isle of Man | 0 | 0.00 | 0 | 0.00 | 0 | 0.00 | 0 | 0.00 |
| Israel | 453 | 3.95 | 936 | 8.16 | 1307 | 11.41 | 2486 | 21.69 |
| Italy | 574 | 0.88 | 1029 | 1.57 | 1750 | 2.68 | 3691 | 5.65 |
| Jamaica | 31 | 0.66 | 223 | 4.71 | 227 | 4.78 | 324 | 6.83 |
| Japan | 822 | 0.64 | 499 | 0.39 | 2677 | 2.09 | 5024 | 3.92 |
| Jersey | 0 | 0.00 | 0 | 0.00 | 0 | 0.00 | 0 | 0.00 |
| Jordan | 214 | 1.37 | 425 | 2.72 | 857 | 5.48 | 1725 | 11.03 |
| Kazakhstan | 128 | 0.12 | 206 | 0.19 | 4137 | 3.86 | 6888 | 6.43 |
| Kenya | 1363 | 1.36 | 3383 | 3.37 | 8023 | 7.99 | 16764 | 16.68 |
| Kiribati | 0 | 0.00 | 0 | 0.00 | 0 | 0.00 | 0 | 0.00 |
| Kosovo | 1 | 0.07 | 0 | 0.00 | 24 | 1.43 | 166 | 10.10 |
| Kuwait | 56 | 1.94 | 130 | 4.48 | 33 | 1.13 | 59 | 2.02 |
| Kyrgyzstan | 42 | 0.22 | 98 | 0.52 | 122 | 0.65 | 352 | 1.87 |
| Lao PDR | 8135 | 7.95 | 10993 | 10.74 | 3481 | 3.40 | 6086 | 5.95 |
| Latvia | 0 | 0.00 | 97 | 1.43 | 127 | 1.87 | 245 | 3.61 |
| Lebanon | 172 | 6.38 | 304 | 11.25 | 432 | 16.02 | 945 | 35.03 |
| Lesotho | 15 | 0.49 | 338 | 10.73 | 0 | 0.00 | 0 | 0.00 |
| Liberia | 594 | 1.88 | 1130 | 3.58 | 3912 | 12.41 | 7688 | 24.38 |
| Libya | 1388 | 0.15 | 15279 | 1.62 | 5390 | 0.57 | 11916 | 1.26 |
| Liechtenstein | 0 | 0.00 | 0 | 0.00 | 0 | 0.00 | 0 | 0.00 |
| Lithuania | 0 | 0.00 | 15 | 0.26 | 117 | 2.06 | 146 | 2.57 |
| Luxembourg | 0 | 0.00 | 0 | 0.00 | 27 | 2.49 | 67 | 6.12 |
| Macao | 0 | 0.00 | 0 | 0.00 | 0 | 0.00 | 0 | 0.00 |
| Macedonia | 0 | 0.00 | 19 | 0.31 | 27 | 0.45 | 27 | 0.45 |
| Madagascar | 1842 | 1.73 | 3266 | 3.06 | 2191 | 2.06 | 2423 | 2.27 |
| Malawi | 333 | 1.48 | 895 | 3.97 | 415 | 1.84 | 690 | 3.06 |
| Malaysia | 697 | 0.48 | 1887 | 1.29 | 2849 | 1.95 | 3904 | 2.67 |
| Maldives | 0 | 0.00 | 0 | 0.00 | 0 | 0.00 | 0 | 0.00 |
| Mali | 4134 | 0.73 | 2991 | 0.53 | 11903 | 2.11 | 25554 | 4.52 |
| Malta | 0 | 0.00 | 0 | 0.00 | 0 | 0.00 | 0 | 0.00 |
| Marshall Islands | 0 | 0.00 | 0 | 0.00 | 0 | 0.00 | 0 | 0.00 |
| Mauritania | 58 | 0.01 | 128 | 0.02 | 7082 | 1.19 | 15476 | 2.60 |
| Mauritius | 0 | 0.00 | 0 | 0.00 | 0 | 0.00 | 0 | 0.00 |
| Mexico | 1995 | 0.39 | 4159 | 0.82 | 29266 | 5.79 | 36231 | 7.17 |
| Moldova | 0 | 0.00 | 0 | 0.00 | 8 | 5.42 | 2 | 1.59 |
| Monaco | 0 | 0.00 | 0 | 0.00 | 0 | 0.00 | 0 | 0.00 |
| Mongolia | 31 | 0.01 | 56 | 0.02 | 6941 | 3.02 | 11988 | 5.22 |
| Montenegro | 4 | 0.09 | 100 | 2.11 | 4 | 0.08 | 29 | 0.62 |
| Montserrat | 0 | 0.00 | 0 | 0.00 | 0 | 0.00 | 0 | 0.00 |
| Morocco | 875 | 0.94 | 15966 | 17.13 | 5017 | 5.38 | 8558 | 9.18 |
| Mozambique | 480 | 0.50 | 1948 | 2.03 | 4912 | 5.12 | 9796 | 10.21 |
| Myanmar | 2124 | 1.02 | 3584 | 1.73 | 4686 | 2.26 | 6038 | 2.91 |
| Namibia | 45 | 0.02 | 227 | 0.08 | 5207 | 1.92 | 9848 | 3.63 |
| Nauru | 0 | 0.00 | 0 | 0.00 | 0 | 0.00 | 0 | 0.00 |
| Nepal | 766 | 1.37 | 1473 | 2.63 | 719 | 1.28 | 1320 | 2.36 |
| Netherlands | 77 | 1.35 | 143 | 2.48 | 404 | 7.02 | 795 | 13.81 |
| New Caledonia | 3 | 0.02 | 13 | 0.10 | 11 | 0.08 | 43 | 0.32 |
| New Zealand | 285 | 0.38 | 994 | 1.31 | 617 | 0.81 | 1311 | 1.72 |
| Nicaragua | 91 | 0.23 | 261 | 0.65 | 3099 | 7.77 | 5474 | 13.73 |
| Niger | 96 | 0.02 | 313 | 0.05 | 7150 | 1.14 | 20097 | 3.21 |
| Nigeria | 2120 | 4.03 | 3714 | 7.05 | 9584 | 18.19 | 19591 | 37.19 |
| Niue | 0 | 0.00 | 0 | 0.00 | 0 | 0.00 | 0 | 0.00 |
| Norfolk Island | 0 | 0.00 | 0 | 0.00 | 0 | 0.00 | 0 | 0.00 |
| Northern Cyprus | 2 | 0.19 | 2 | 0.28 | 21 | 2.46 | 41 | 4.69 |
| Northern Mariana Islands | 0 | 0.00 | 0 | 0.00 | 0 | 0.00 | 0 | 0.00 |
| Norway | 99 | 0.10 | 201 | 0.19 | 277 | 0.27 | 1236 | 1.19 |
| Oman | 117 | 0.08 | 199 | 0.14 | 1517 | 1.05 | 3093 | 2.15 |
| Pakistan | 1042 | 0.69 | 1406 | 0.93 | 8423 | 5.55 | 15031 | 9.90 |
| Palau | 0 | 0.00 | 0 | 0.00 | 0 | 0.00 | 0 | 0.00 |
| Palestine | 61 | 3.56 | 113 | 6.57 | 189 | 10.98 | 385 | 22.36 |
| Panama | 667 | 1.75 | 648 | 1.70 | 2124 | 5.58 | 3437 | 9.04 |
| Papua New Guinea | 54 | 0.02 | 119 | 0.05 | 1264 | 0.53 | 1917 | 0.80 |
| Paraguay | 295 | 0.35 | 1170 | 1.40 | 6801 | 8.16 | 12337 | 14.79 |
| Peru | 433 | 0.07 | 1228 | 0.18 | 13215 | 1.99 | 18545 | 2.79 |
| Philippines | 654 | 0.72 | 1917 | 2.12 | 4851 | 5.37 | 8365 | 9.26 |
| Pitcairn Islands | 0 | 0.00 | 0 | 0.00 | 0 | 0.00 | 0 | 0.00 |
| Poland | 87 | 0.11 | 101 | 0.12 | 969 | 1.19 | 2139 | 2.63 |
| Portugal | 75 | 0.54 | 171 | 1.22 | 437 | 3.11 | 748 | 5.33 |
| Puerto Rico | 64 | 1.94 | 172 | 5.23 | 0 | 0.00 | 0 | 0.00 |
| Qatar | 16 | 0.94 | 34 | 2.00 | 46 | 2.67 | 116 | 6.74 |
| Republic of Congo | 637 | 0.49 | 805 | 0.62 | 857 | 0.66 | 929 | 0.71 |
| Republic of Korea | 284 | 0.99 | 114 | 0.40 | 1592 | 5.56 | 1687 | 5.89 |
| Romania | 75 | 0.20 | 62 | 0.17 | 1180 | 3.15 | 1569 | 4.19 |
| Russian Federation | 243 | 0.00 | 1332 | 0.01 | 19824 | 0.21 | 44323 | 0.47 |
| Rwanda | 108 | 3.33 | 339 | 10.41 | 407 | 12.51 | 694 | 21.32 |
| Saint Helena | 0 | 0.00 | 0 | 0.00 | 0 | 0.00 | 0 | 0.00 |
| Saint Kitts and Nevis | 1 | 1.44 | 7 | 6.83 | 0 | 0.00 | 0 | 0.00 |
| Saint Lucia | 1 | 0.34 | 0 | 0.00 | 3 | 0.84 | 5 | 1.53 |
| Saint Pierre and Miquelon | 0 | 0.00 | 0 | 0.00 | 0 | 0.00 | 0 | 0.00 |
| Saint Vincent and the Grenadines | 0 | 0.00 | 0 | 0.00 | 0 | 0.00 | 0 | 0.00 |
| Saint-Bartholemy | 0 | 0.00 | 0 | 0.00 | 0 | 0.00 | 0 | 0.00 |
| Saint-Martin | 0 | 0.00 | 0 | 0.00 | 0 | 0.00 | 0 | 0.00 |
| Samoa | 349 | 25.24 | 403 | 29.10 | 1 | 0.06 | 6 | 0.43 |
| San Marino | 0 | 0.00 | 0 | 0.00 | 0 | 0.00 | 0 | 0.00 |
| Sao Tome and Principe | 4 | 2.09 | 11 | 5.45 | 5 | 2.24 | 11 | 5.50 |
| Saudi Arabia | 547 | 0.71 | 1123 | 1.46 | 3639 | 4.73 | 6930 | 9.00 |
| Scarborough Reef | 0 | 0.00 | 0 | 0.00 | 0 | 0.00 | 0 | 0.00 |
| Senegal | 890 | 1.85 | 900 | 1.87 | 4000 | 8.30 | 8574 | 17.80 |
| Serbia | 4 | 0.06 | 20 | 0.31 | 314 | 4.95 | 676 | 10.66 |
| Serranilla Bank | 0 | 0.00 | 0 | 0.00 | 0 | 0.00 | 0 | 0.00 |
| Seychelles | 0 | 0.00 | 0 | 0.00 | 0 | 0.00 | 0 | 0.00 |
| Siachen Glacier | 1 | 0.23 | 2 | 0.39 | 3 | 0.54 | 3 | 0.61 |
| Sierra Leone | 494 | 5.34 | 557 | 6.02 | 1157 | 12.52 | 539 | 5.84 |
| Singapore | 0 | 0.00 | 0 | 0.00 | 0 | 0.00 | 0 | 0.00 |
| Sint Maarten | 0 | 0.00 | 0 | 0.00 | 0 | 0.00 | 0 | 0.00 |
| Slovakia | 46 | 0.40 | 64 | 0.55 | 38 | 0.33 | 59 | 0.50 |
| Slovenia | 60 | 0.50 | 79 | 0.66 | 100 | 0.83 | 103 | 0.86 |
| Solomon Islands | 30 | 0.20 | 46 | 0.31 | 15 | 0.10 | 45 | 0.31 |
| Somalia | 162 | 0.49 | 313 | 0.94 | 598 | 1.79 | 2692 | 8.07 |
| Somaliland | 14 | 0.06 | 121 | 0.54 | 1936 | 8.72 | 3526 | 15.87 |
| South Africa | 815 | 0.69 | 4878 | 4.14 | 6886 | 5.84 | 9193 | 7.79 |
| South Georgia and South Sandwich Islands | 0 | 0.00 | 0 | 0.00 | 0 | 0.00 | 0 | 0.00 |
| South Sudan | 1198 | 2.58 | 2441 | 5.26 | 4929 | 10.63 | 11326 | 24.42 |
| Spain | 235 | 0.18 | 597 | 0.46 | 4981 | 3.80 | 9181 | 7.01 |
| Spratly Islands | 0 | 0.00 | 0 | 0.00 | 0 | 0.00 | 0 | 0.00 |
| Sri Lanka | 265 | 1.00 | 2185 | 8.27 | 1343 | 5.08 | 1942 | 7.35 |
| Sudan | 452 | 0.26 | 2364 | 1.36 | 8684 | 5.00 | 18072 | 10.41 |
| Suriname | 22 | 0.02 | 52 | 0.04 | 709 | 0.58 | 1135 | 0.93 |
| Swaziland | 0 | 0.00 | 5 | 0.14 | 1 | 0.04 | 29 | 0.84 |
| Sweden | 31 | 0.03 | 61 | 0.07 | 507 | 0.56 | 601 | 0.67 |
| Switzerland | 3 | 0.07 | 0 | 0.00 | 27 | 0.52 | 34 | 0.66 |
| Syria | 121 | 2.01 | 202 | 3.36 | 452 | 7.51 | 547 | 9.10 |
| Taiwan | 233 | 1.38 | 442 | 2.62 | 354 | 2.10 | 545 | 3.23 |
| Tajikistan | 21 | 0.05 | 42 | 0.10 | 672 | 1.63 | 1321 | 3.21 |
| Tanzania | 2114 | 0.72 | 6843 | 2.34 | 20123 | 6.87 | 24259 | 8.28 |
| Thailand | 1351 | 0.81 | 2280 | 1.37 | 10991 | 6.62 | 21178 | 12.75 |
| The Gambia | 86 | 2.76 | 60 | 1.93 | 271 | 8.67 | 484 | 15.51 |
| Timor-Leste | 187 | 4.75 | 749 | 19.01 | 399 | 10.11 | 1226 | 31.09 |
| Togo | 196 | 6.03 | 157 | 4.82 | 236 | 7.27 | 634 | 19.50 |
| Tonga | 0 | 0.00 | 0 | 0.00 | 0 | 0.00 | 0 | 0.00 |
| Trinidad and Tobago | 0 | 0.00 | 219 | 5.77 | 47 | 1.23 | 122 | 3.20 |
| Tunisia | 81 | 1.04 | 855 | 10.94 | 236 | 3.02 | 433 | 5.54 |
| Turkey | 859 | 0.67 | 1689 | 1.31 | 17895 | 13.87 | 34350 | 26.63 |
| Turkmenistan | 88 | 0.30 | 144 | 0.49 | 452 | 1.53 | 1001 | 3.40 |
| Turks and Caicos Islands | 3 | 1.90 | 5 | 2.87 | 3 | 1.89 | 5 | 2.86 |
| Tuvalu | 0 | 0.00 | 0 | 0.00 | 0 | 0.00 | 0 | 0.00 |
| Uganda | 762 | 2.89 | 2355 | 8.93 | 3530 | 13.38 | 6846 | 25.95 |
| Ukraine | 35 | 0.12 | 201 | 0.71 | 8845 | 31.13 | 13118 | 46.16 |
| United Arab Emirates | 267 | 1.25 | 568 | 2.66 | 716 | 3.35 | 1582 | 7.40 |
| United Kingdom | 42 | 0.18 | 184 | 0.80 | 456 | 1.98 | 577 | 2.50 |
| United States | 5173 | 0.22 | 13186 | 0.55 | 27741 | 1.16 | 46365 | 1.94 |
| United States Minor Outlying Islands | 0 | 0.00 | 0 | 0.00 | 0 | 0.00 | 0 | 0.00 |
| United States Virgin Islands | 0 | 0.00 | 0 | 0.00 | 0 | 0.00 | 0 | 0.00 |
| Uruguay | 195 | 2.35 | 230 | 2.76 | 3059 | 36.77 | 3561 | 42.80 |
| US Naval Base Guantanamo Bay | 0 | 0.00 | 0 | 0.00 | 0 | 0.00 | 0 | 0.00 |
| Uzbekistan | 102 | 0.26 | 259 | 0.65 | 1130 | 2.83 | 2645 | 6.63 |
| Vanuatu | 35 | 1.02 | 48 | 1.37 | 2 | 0.05 | 199 | 5.72 |
| Venezuela | 1150 | 0.21 | 2611 | 0.48 | 16547 | 3.06 | 27879 | 5.16 |
| Vietnam | 1669 | 1.27 | 2405 | 1.83 | 8850 | 6.74 | 17848 | 13.60 |
| Wallis and Futuna Islands | 0 | 0.00 | 0 | 0.00 | 0 | 0.00 | 0 | 0.00 |
| Western Sahara | 9 | 0.04 | 25 | 0.12 | 268 | 1.24 | 529 | 2.45 |
| Yemen | 240 | 0.31 | 419 | 0.54 | 1260 | 1.61 | 2944 | 3.76 |
| Zambia | 2622 | 0.94 | 5623 | 2.02 | 2580 | 0.93 | 6108 | 2.19 |
| Zimbabwe | 357 | 0.46 | 1341 | 1.72 | 4420 | 5.66 | 7345 | 9.41 |

| ***Extended Data Table 4.*** *The area and proportion of intact land requiring conservation within each country that is at risk of habitat conversion by 2030 and 2050 based on Shared Socioeconomic Pathways 3 (pessimistic) and "ensemble (an average of SSP1, 2, and 3).* | | | | | | | | |
| --- | --- | --- | --- | --- | --- | --- | --- | --- |
| **Country** | **Habitat loss by 2030 (SSP3) (km2)** | **Habitat loss by 2030 (SSP3) (%)** | **Habitat loss by 2050 (SSP3) (km2)** | **Habitat loss by 2050 (SSP3) (%)** | **Ensemble 2030 (km2)** | **Ensemble 2030 (%)** | **Ensemble2050 (km2)** | **Ensemble 2050 (%)** |
| Afghanistan | 6383 | 11.04 | 9458 | 16.36 | 2244 | 3.88 | 3317 | 5.74 |
| Akrotiri | 0 | 0.00 | 0 | 0.00 | 0 | 0.00 | 0 | 0.00 |
| Aland Islands | 0 | 0.00 | 0 | 0.00 | 0 | 0.00 | 0 | 0.00 |
| Albania | 34 | 0.66 | 49 | 0.94 | 35 | 0.67 | 94 | 1.80 |
| Algeria | 15942 | 1.32 | 28298 | 2.35 | 7807 | 0.65 | 14887 | 1.23 |
| American Samoa | 0 | 0.00 | 0 | 0.00 | 0 | 0.00 | 0 | 0.00 |
| Andorra | 10 | 1.49 | 30 | 4.62 | 0 | 0.00 | 0 | 0.00 |
| Angola | 38729 | 27.68 | 63207 | 45.17 | 14545 | 10.39 | 24334 | 17.39 |
| Anguilla | 0 | 0.00 | 0 | 0.00 | 0 | 0.00 | 0 | 0.00 |
| Antigua and Barbuda | 0 | 0.00 | 0 | 0.00 | 0 | 0.00 | 0 | 0.00 |
| Argentina | 5157 | 1.03 | 12617 | 2.52 | 10799 | 2.16 | 21463 | 4.30 |
| Armenia | 18 | 0.27 | 16 | 0.24 | 227 | 3.45 | 511 | 7.78 |
| Aruba | 0 | 0.00 | 0 | 0.00 | 0 | 0.00 | 0 | 0.00 |
| Ashmore and Cartier Islands | 0 | 0.00 | 0 | 0.00 | 0 | 0.00 | 0 | 0.00 |
| Australia | 18390 | 0.72 | 20722 | 0.81 | 20972 | 0.82 | 39962 | 1.57 |
| Austria | 419 | 2.48 | 626 | 3.71 | 0 | 0.00 | 0 | 0.00 |
| Azerbaijan | 181 | 1.22 | 86 | 0.58 | 686 | 4.64 | 1394 | 9.43 |
| Bahamas | 9 | 0.23 | 16 | 0.40 | 125 | 3.11 | 190 | 4.75 |
| Bahrain | 10 | 2.30 | 19 | 4.19 | 0 | 0.00 | 0 | 0.00 |
| Baikonur Cosmodrome | 0 | 0.00 | 0 | 0.00 | 0 | 0.00 | 0 | 0.00 |
| Bangladesh | 1866 | 12.44 | 3083 | 20.56 | 896 | 5.97 | 1444 | 9.63 |
| Barbados | 13 | 6.23 | 32 | 16.00 | 0 | 0.00 | 0 | 0.00 |
| Belarus | 39 | 0.20 | 15 | 0.08 | 207 | 1.03 | 355 | 1.77 |
| Belgium | 312 | 5.36 | 605 | 10.38 | 0 | 0.00 | 0 | 0.00 |
| Belize | 11 | 0.06 | 20 | 0.11 | 215 | 1.21 | 296 | 1.66 |
| Benin | 5863 | 21.91 | 11636 | 43.49 | 3648 | 13.64 | 7467 | 27.91 |
| Bermuda | 0 | 0.00 | 0 | 0.00 | 0 | 0.00 | 0 | 0.00 |
| Bhutan | 1891 | 9.27 | 2454 | 12.03 | 781 | 3.82 | 1063 | 5.21 |
| Bolivia | 6534 | 1.65 | 11589 | 2.92 | 8533 | 2.15 | 14186 | 3.58 |
| Bosnia and Herzegovina | 5 | 0.12 | 3 | 0.08 | 54 | 1.36 | 75 | 1.88 |
| Botswana | 32464 | 21.36 | 40687 | 26.77 | 11739 | 7.72 | 15156 | 9.97 |
| Brazil | 70562 | 2.05 | 121190 | 3.53 | 64783 | 1.88 | 110387 | 3.21 |
| British Indian Ocean Territory | 0 | 0.00 | 0 | 0.00 | 0 | 0.00 | 0 | 0.00 |
| British Virgin Islands | 0 | 0.00 | 0 | 0.00 | 0 | 0.00 | 0 | 0.00 |
| Brunei Darussalam | 564 | 11.58 | 1078 | 22.12 | 206 | 4.23 | 393 | 8.08 |
| Bulgaria | 192 | 0.76 | 482 | 1.91 | 0 | 0.00 | 0 | 0.00 |
| Burkina Faso | 5977 | 19.15 | 14134 | 45.29 | 0 | 0.00 | 0 | 0.00 |
| Burundi | 384 | 22.11 | 685 | 39.44 | 178 | 10.24 | 294 | 16.93 |
| Cambodia | 471 | 0.69 | 714 | 1.05 | 1560 | 2.29 | 3128 | 4.59 |
| Cameroon | 17335 | 13.66 | 40810 | 32.15 | 8368 | 6.59 | 18870 | 14.87 |
| Canada | 31374 | 0.44 | 39111 | 0.55 | 19063 | 0.27 | 28290 | 0.40 |
| Cape Verde | 17 | 1.22 | 89 | 6.36 | 0 | 0.00 | 0 | 0.00 |
| Cayman Islands | 0 | 0.00 | 0 | 0.00 | 0 | 0.00 | 0 | 0.00 |
| Central African Republic | 9198 | 3.98 | 23845 | 10.33 | 4392 | 1.90 | 10520 | 4.56 |
| Chad | 15705 | 2.95 | 36462 | 6.84 | 11086 | 2.08 | 24000 | 4.50 |
| Chile | 2492 | 0.86 | 6357 | 2.20 | 3913 | 1.35 | 6319 | 2.19 |
| China | 2388 | 0.16 | 2059 | 0.13 | 14282 | 0.93 | 23081 | 1.50 |
| Clipperton Island | 0 | 0.00 | 0 | 0.00 | 0 | 0.00 | 0 | 0.00 |
| Colombia | 1060 | 0.20 | 3008 | 0.58 | 5841 | 1.12 | 9669 | 1.85 |
| Comoros | 0 | 86.16 | 0 | 35.90 | 0 | 0.00 | 0 | 0.00 |
| Cook Islands | 0 | 0.00 | 0 | 0.00 | 0 | 0.00 | 0 | 0.00 |
| Coral Sea Islands | 0 | 0.00 | 0 | 0.00 | 0 | 0.00 | 0 | 0.00 |
| Costa Rica | 196 | 0.89 | 682 | 3.09 | 381 | 1.73 | 754 | 3.42 |
| Cote d'Ivoire | 14385 | 36.78 | 18877 | 48.26 | 5788 | 14.80 | 8181 | 20.92 |
| Croatia | 20 | 0.11 | 56 | 0.32 | 268 | 1.54 | 456 | 2.62 |
| Cuba | 8 | 0.03 | 4 | 0.02 | 1173 | 4.84 | 1832 | 7.56 |
| Curacao | 0 | 0.00 | 0 | 0.00 | 0 | 0.00 | 0 | 0.00 |
| Cyprus | 8 | 0.18 | 16 | 0.39 | 41 | 0.97 | 78 | 1.84 |
| Cyprus U.N. Buffer Zone | 0 | 0.00 | 0 | 0.00 | 0 | 0.00 | 0 | 0.00 |
| Czech Republic | 47 | 0.63 | 48 | 0.63 | 0 | 0.00 | 0 | 0.00 |
| Dem. Rep. Korea | 47 | 0.29 | 67 | 0.42 | 213 | 1.33 | 284 | 1.78 |
| Democratic Republic of the Congo | 38301 | 6.95 | 116567 | 21.14 | 18527 | 3.36 | 48126 | 8.73 |
| Denmark | 19 | 0.74 | 59 | 2.36 | 57 | 2.29 | 65 | 2.61 |
| Dhekelia | 0 | 0.00 | 0 | 0.00 | 0 | 0.00 | 0 | 0.00 |
| Djibouti | 46 | 3.27 | 167 | 11.95 | 47 | 3.38 | 128 | 9.15 |
| Dominica | 0 | 0.00 | 0 | 0.00 | 0 | 0.00 | 0 | 0.00 |
| Dominican Republic | 147 | 0.91 | 83 | 0.52 | 1003 | 6.24 | 1496 | 9.31 |
| Ecuador | 7853 | 5.33 | 13069 | 8.87 | 4607 | 3.13 | 7974 | 5.41 |
| Egypt | 10015 | 1.70 | 16132 | 2.73 | 5924 | 1.00 | 13332 | 2.26 |
| El Salvador | 52 | 2.36 | 99 | 4.47 | 157 | 7.05 | 262 | 11.77 |
| Equatorial Guinea | 547 | 9.24 | 1337 | 22.57 | 202 | 3.40 | 472 | 7.97 |
| Eritrea | 3012 | 29.25 | 5753 | 55.86 | 1749 | 16.98 | 3338 | 32.41 |
| Estonia | 5 | 0.08 | 19 | 0.32 | 0 | 0.00 | 0 | 0.00 |
| Ethiopia | 69588 | 24.67 | 151404 | 53.66 | 32038 | 11.36 | 68226 | 24.18 |
| Faeroe Islands | 0 | 65.24 | 0 | 65.24 | 0 | 0.00 | 0 | 0.00 |
| Falkland Islands | 0 | 0.00 | 0 | 0.00 | 1 | 0.02 | 0 | 0.01 |
| Federated States of Micronesia | 0 | 0.00 | 0 | 0.00 | 0 | 0.00 | 0 | 0.00 |
| Fiji | 0 | 0.00 | 0 | 0.00 | 0 | 0.00 | 0 | 0.00 |
| Finland | 22 | 0.03 | 46 | 0.07 | 203 | 0.31 | 269 | 0.41 |
| France | 4376 | 5.17 | 8294 | 9.80 | 2242 | 2.65 | 4379 | 5.17 |
| French Polynesia | 0 | 0.00 | 0 | 0.00 | 0 | 0.00 | 0 | 0.00 |
| French Southern and Antarctic Lands | 0 | 0.00 | 0 | 0.00 | 0 | 0.00 | 0 | 0.00 |
| Gabon | 10555 | 14.79 | 29360 | 41.15 | 3778 | 5.29 | 10010 | 14.03 |
| Georgia | 92 | 0.42 | 51 | 0.24 | 381 | 1.75 | 1093 | 5.01 |
| Germany | 1208 | 1.54 | 1461 | 1.86 | 1651 | 2.11 | 3158 | 4.03 |
| Ghana | 11465 | 50.33 | 15095 | 66.27 | 5008 | 21.99 | 6983 | 30.66 |
| Gibraltar | 0 | 0.00 | 0 | 0.00 | 0 | 0.00 | 0 | 0.00 |
| Greece | 444 | 2.33 | 1180 | 6.19 | 483 | 2.53 | 1121 | 5.88 |
| Greenland | 1 | 0.00 | 2 | 0.00 | 92 | 0.04 | 287 | 0.12 |
| Grenada | 0 | 0.00 | 0 | 0.00 | 0 | 0.00 | 0 | 0.00 |
| Guam | 0 | 0.00 | 0 | 0.00 | 0 | 0.00 | 0 | 0.00 |
| Guatemala | 9668 | 18.90 | 14047 | 27.46 | 4551 | 8.90 | 7546 | 14.75 |
| Guernsey | 0 | 0.00 | 0 | 0.00 | 0 | 0.00 | 0 | 0.00 |
| Guinea | 22856 | 47.80 | 32412 | 67.79 | 11550 | 24.16 | 17736 | 37.10 |
| Guinea-Bissau | 2545 | 42.63 | 3699 | 61.97 | 1090 | 18.26 | 1697 | 28.43 |
| Guyana | 1102 | 1.01 | 1066 | 0.98 | 1072 | 0.98 | 1291 | 1.18 |
| Haiti | 113 | 1.77 | 203 | 3.19 | 313 | 4.90 | 322 | 5.04 |
| Heard I. and McDonald Islands | 0 | 0.00 | 0 | 0.00 | 0 | 0.00 | 0 | 0.00 |
| Honduras | 176 | 0.34 | 215 | 0.42 | 1653 | 3.21 | 2931 | 5.69 |
| Hong Kong | 0 | 0.00 | 0 | 0.00 | 0 | 0.00 | 0 | 0.00 |
| Hungary | 64 | 0.73 | 0 | 0.00 | 0 | 0.00 | 0 | 0.00 |
| Iceland | 4 | 0.01 | 6 | 0.02 | 31 | 0.08 | 48 | 0.13 |
| India | 48571 | 11.51 | 73175 | 17.34 | 25418 | 6.02 | 40302 | 9.55 |
| Indian Ocean Territories | 0 | 0.00 | 0 | 0.00 | 0 | 0.00 | 0 | 0.00 |
| Indonesia | 18898 | 3.04 | 27692 | 4.45 | 12310 | 1.98 | 20816 | 3.35 |
| Iran | 5387 | 2.24 | 4573 | 1.90 | 7110 | 2.96 | 11132 | 4.63 |
| Iraq | 1658 | 3.01 | 2036 | 3.70 | 1750 | 3.18 | 2878 | 5.23 |
| Ireland | 491 | 8.72 | 2133 | 37.85 | 217 | 3.85 | 757 | 13.43 |
| Isle of Man | 0 | 0.00 | 0 | 0.00 | 0 | 0.00 | 0 | 0.00 |
| Israel | 2973 | 25.94 | 6037 | 52.66 | 1578 | 13.76 | 3153 | 27.50 |
| Italy | 1324 | 2.03 | 2292 | 3.51 | 1216 | 1.86 | 2337 | 3.58 |
| Jamaica | 38 | 0.80 | 83 | 1.75 | 99 | 2.08 | 210 | 4.43 |
| Japan | 209 | 0.16 | 243 | 0.19 | 1236 | 0.96 | 1922 | 1.50 |
| Jersey | 0 | 0.00 | 0 | 0.00 | 0 | 0.00 | 0 | 0.00 |
| Jordan | 3129 | 20.01 | 5405 | 34.56 | 1400 | 8.95 | 2518 | 16.10 |
| Kazakhstan | 4462 | 4.17 | 3909 | 3.65 | 2909 | 2.72 | 3667 | 3.43 |
| Kenya | 26765 | 26.64 | 34547 | 34.38 | 12050 | 11.99 | 18231 | 18.14 |
| Kiribati | 0 | 0.00 | 0 | 0.00 | 0 | 0.00 | 0 | 0.00 |
| Kosovo | 7 | 0.44 | 16 | 0.98 | 11 | 0.65 | 61 | 3.69 |
| Kuwait | 90 | 3.09 | 122 | 4.18 | 60 | 2.05 | 104 | 3.56 |
| Kyrgyzstan | 338 | 1.80 | 326 | 1.73 | 168 | 0.89 | 258 | 1.37 |
| Lao PDR | 1461 | 1.43 | 1523 | 1.49 | 4359 | 4.26 | 6201 | 6.06 |
| Latvia | 13 | 0.19 | 63 | 0.93 | 0 | 0.00 | 0 | 0.00 |
| Lebanon | 1574 | 58.33 | 1720 | 63.76 | 726 | 26.91 | 990 | 36.68 |
| Lesotho | 1352 | 42.88 | 1667 | 52.88 | 456 | 14.46 | 668 | 21.20 |
| Liberia | 12195 | 38.67 | 24343 | 77.19 | 5567 | 17.65 | 11054 | 35.05 |
| Libya | 13225 | 1.40 | 19866 | 2.10 | 6668 | 0.71 | 15687 | 1.66 |
| Liechtenstein | 0 | 0.00 | 0 | 0.00 | 0 | 0.00 | 0 | 0.00 |
| Lithuania | 54 | 0.96 | 175 | 3.08 | 57 | 1.01 | 112 | 1.97 |
| Luxembourg | 44 | 3.98 | 41 | 3.75 | 0 | 0.00 | 0 | 0.00 |
| Macao | 0 | 0.00 | 0 | 0.00 | 0 | 0.00 | 0 | 0.00 |
| Macedonia | 69 | 1.13 | 180 | 2.97 | 32 | 0.53 | 75 | 1.24 |
| Madagascar | 22226 | 20.86 | 32800 | 30.78 | 8753 | 8.21 | 12829 | 12.04 |
| Malawi | 7346 | 32.60 | 13023 | 57.81 | 2698 | 11.97 | 4869 | 21.61 |
| Malaysia | 10780 | 7.37 | 19279 | 13.18 | 4775 | 3.27 | 8357 | 5.71 |
| Maldives | 0 | 0.00 | 0 | 0.00 | 0 | 0.00 | 0 | 0.00 |
| Mali | 34136 | 6.04 | 64902 | 11.49 | 16724 | 2.96 | 31149 | 5.52 |
| Malta | 1 | 0.43 | 1 | 0.44 | 0 | 0.00 | 0 | 0.00 |
| Marshall Islands | 0 | 0.00 | 0 | 0.00 | 0 | 0.00 | 0 | 0.00 |
| Mauritania | 14856 | 2.50 | 19689 | 3.31 | 7332 | 1.23 | 11765 | 1.98 |
| Mauritius | 0 | 0.00 | 0 | 0.00 | 0 | 0.00 | 0 | 0.00 |
| Mexico | 3659 | 0.72 | 5469 | 1.08 | 11640 | 2.30 | 15286 | 3.02 |
| Moldova | 0 | 0.00 | 0 | 0.00 | 0 | 0.00 | 0 | 0.00 |
| Monaco | 0 | 0.00 | 0 | 0.00 | 0 | 0.00 | 0 | 0.00 |
| Mongolia | 12828 | 5.59 | 12316 | 5.37 | 6600 | 2.88 | 8120 | 3.54 |
| Montenegro | 63 | 1.34 | 68 | 1.43 | 24 | 0.50 | 66 | 1.39 |
| Montserrat | 0 | 0.00 | 0 | 0.00 | 0 | 0.00 | 0 | 0.00 |
| Morocco | 2678 | 2.87 | 3452 | 3.70 | 2857 | 3.06 | 9326 | 10.00 |
| Mozambique | 23983 | 25.00 | 34215 | 35.67 | 9792 | 10.21 | 15320 | 15.97 |
| Myanmar | 5669 | 2.73 | 6275 | 3.02 | 4160 | 2.00 | 5299 | 2.55 |
| Namibia | 36088 | 13.32 | 40024 | 14.77 | 13780 | 5.09 | 16700 | 6.16 |
| Nauru | 0 | 0.00 | 0 | 0.00 | 0 | 0.00 | 0 | 0.00 |
| Nepal | 1688 | 3.02 | 2776 | 4.96 | 1057 | 1.89 | 1856 | 3.32 |
| Netherlands | 196 | 3.40 | 384 | 6.68 | 226 | 3.92 | 440 | 7.65 |
| New Caledonia | 631 | 4.58 | 677 | 4.92 | 215 | 1.56 | 244 | 1.78 |
| New Zealand | 6674 | 8.78 | 9788 | 12.88 | 2526 | 3.32 | 4031 | 5.30 |
| Nicaragua | 97 | 0.24 | 100 | 0.25 | 1096 | 2.75 | 1945 | 4.88 |
| Niger | 11579 | 1.85 | 25757 | 4.12 | 0 | 0.00 | 0 | 0.00 |
| Nigeria | 11392 | 21.63 | 21928 | 41.63 | 7699 | 14.62 | 15078 | 28.62 |
| Niue | 0 | 0.00 | 0 | 0.00 | 0 | 0.00 | 0 | 0.00 |
| Norfolk Island | 0 | 0.00 | 0 | 0.00 | 0 | 0.00 | 0 | 0.00 |
| Northern Cyprus | 2 | 0.18 | 3 | 0.39 | 8 | 0.94 | 16 | 1.79 |
| Northern Mariana Islands | 0 | 0.00 | 0 | 0.00 | 0 | 0.00 | 0 | 0.00 |
| Norway | 2246 | 2.16 | 4815 | 4.62 | 874 | 0.84 | 2084 | 2.00 |
| Oman | 2959 | 2.06 | 2670 | 1.86 | 1531 | 1.06 | 1987 | 1.38 |
| Pakistan | 8259 | 5.44 | 16685 | 10.99 | 5908 | 3.89 | 11041 | 7.27 |
| Palau | 0 | 0.00 | 0 | 0.00 | 0 | 0.00 | 0 | 0.00 |
| Palestine | 354 | 20.61 | 756 | 43.93 | 202 | 11.72 | 418 | 24.29 |
| Panama | 247 | 0.65 | 878 | 2.31 | 1013 | 2.66 | 1654 | 4.35 |
| Papua New Guinea | 5304 | 2.21 | 6619 | 2.75 | 2207 | 0.92 | 2885 | 1.20 |
| Paraguay | 1217 | 1.46 | 1454 | 1.74 | 2771 | 3.32 | 4987 | 5.98 |
| Peru | 30721 | 4.62 | 76572 | 11.53 | 14790 | 2.23 | 32115 | 4.83 |
| Philippines | 1145 | 1.27 | 2712 | 3.00 | 2216 | 2.45 | 4332 | 4.79 |
| Pitcairn Islands | 0 | 0.00 | 0 | 0.00 | 0 | 0.00 | 0 | 0.00 |
| Poland | 923 | 1.14 | 1252 | 1.54 | 660 | 0.81 | 1164 | 1.43 |
| Portugal | 90 | 0.64 | 115 | 0.82 | 201 | 1.43 | 345 | 2.45 |
| Puerto Rico | 445 | 13.49 | 768 | 23.30 | 170 | 5.14 | 314 | 9.51 |
| Qatar | 47 | 2.73 | 64 | 3.72 | 0 | 0.00 | 0 | 0.00 |
| Republic of Congo | 14265 | 10.98 | 33929 | 26.10 | 5253 | 4.04 | 11888 | 9.15 |
| Republic of Korea | 1349 | 4.71 | 1905 | 6.66 | 1075 | 3.76 | 1235 | 4.32 |
| Romania | 872 | 2.33 | 835 | 2.23 | 709 | 1.89 | 822 | 2.20 |
| Russian Federation | 2204 | 0.02 | 2119 | 0.02 | 7424 | 0.08 | 15925 | 0.17 |
| Rwanda | 1093 | 33.57 | 1687 | 51.81 | 536 | 16.47 | 906 | 27.84 |
| Saint Helena | 0 | 0.00 | 0 | 0.00 | 0 | 0.00 | 0 | 0.00 |
| Saint Kitts and Nevis | 6 | 6.20 | 11 | 11.04 | 0 | 0.00 | 0 | 0.00 |
| Saint Lucia | 3 | 1.00 | 7 | 2.06 | 0 | 0.00 | 0 | 0.00 |
| Saint Pierre and Miquelon | 0 | 0.00 | 0 | 0.00 | 0 | 0.00 | 0 | 0.00 |
| Saint Vincent and the Grenadines | 0 | 0.00 | 0 | 0.00 | 0 | 0.00 | 0 | 0.00 |
| Saint-Bartholemy | 0 | 0.00 | 0 | 0.00 | 0 | 0.00 | 0 | 0.00 |
| Saint-Martin | 0 | 0.00 | 0 | 0.00 | 0 | 0.00 | 0 | 0.00 |
| Samoa | 1 | 0.05 | 3 | 0.22 | 117 | 8.45 | 137 | 9.92 |
| San Marino | 0 | 0.00 | 0 | 0.00 | 0 | 0.00 | 0 | 0.00 |
| Sao Tome and Principe | 0 | 0.10 | 7 | 3.38 | 3 | 1.48 | 10 | 4.77 |
| Saudi Arabia | 3661 | 4.75 | 6465 | 8.40 | 2616 | 3.40 | 4840 | 6.28 |
| Scarborough Reef | 0 | 0.00 | 0 | 0.00 | 0 | 0.00 | 0 | 0.00 |
| Senegal | 13731 | 28.50 | 20338 | 42.22 | 6207 | 12.88 | 9938 | 20.63 |
| Serbia | 47 | 0.75 | 51 | 0.80 | 0 | 0.00 | 0 | 0.00 |
| Serranilla Bank | 0 | 0.00 | 0 | 0.00 | 0 | 0.00 | 0 | 0.00 |
| Seychelles | 0 | 0.00 | 0 | 0.00 | 0 | 0.00 | 0 | 0.00 |
| Siachen Glacier | 0 | 0.00 | 0 | 0.00 | 0 | 0.00 | 0 | 0.00 |
| Sierra Leone | 6373 | 68.95 | 9243 | 100.00 | 2675 | 28.94 | 3446 | 37.29 |
| Singapore | 0 | 0.00 | 0 | 0.00 | 0 | 0.00 | 0 | 0.00 |
| Sint Maarten | 0 | 0.00 | 0 | 0.00 | 0 | 0.00 | 0 | 0.00 |
| Slovakia | 110 | 0.94 | 24 | 0.21 | 0 | 0.00 | 0 | 0.00 |
| Slovenia | 229 | 1.91 | 188 | 1.57 | 0 | 0.00 | 0 | 0.00 |
| Solomon Islands | 18 | 0.12 | 1966 | 13.34 | 21 | 0.14 | 686 | 4.65 |
| Somalia | 4037 | 12.11 | 5713 | 17.13 | 1599 | 4.80 | 2906 | 8.72 |
| Somaliland | 1374 | 6.18 | 2635 | 11.86 | 1108 | 4.99 | 2094 | 9.43 |
| South Africa | 44889 | 38.06 | 51588 | 43.74 | 17530 | 14.86 | 21886 | 18.56 |
| South Georgia and South Sandwich Islands | 0 | 0.00 | 0 | 0.00 | 0 | 0.00 | 0 | 0.00 |
| South Sudan | 2139 | 4.61 | 3201 | 6.90 | 2755 | 5.94 | 5656 | 12.20 |
| Spain | 1541 | 1.18 | 4659 | 3.56 | 2252 | 1.72 | 4812 | 3.67 |
| Spratly Islands | 0 | 0.00 | 0 | 0.00 | 0 | 0.00 | 0 | 0.00 |
| Sri Lanka | 1729 | 6.54 | 1675 | 6.34 | 1112 | 4.21 | 1934 | 7.32 |
| Sudan | 5143 | 2.96 | 10767 | 6.20 | 4760 | 2.74 | 10401 | 5.99 |
| Suriname | 70 | 0.06 | 137 | 0.11 | 267 | 0.22 | 441 | 0.36 |
| Swaziland | 2230 | 65.39 | 2885 | 84.60 | 744 | 21.81 | 973 | 28.53 |
| Sweden | 91 | 0.10 | 169 | 0.19 | 210 | 0.23 | 277 | 0.31 |
| Switzerland | 125 | 2.45 | 332 | 6.49 | 0 | 0.00 | 0 | 0.00 |
| Syria | 1061 | 17.63 | 1148 | 19.09 | 545 | 9.05 | 633 | 10.52 |
| Taiwan | 1614 | 9.57 | 1483 | 8.79 | 734 | 4.35 | 823 | 4.88 |
| Tajikistan | 312 | 0.76 | 374 | 0.91 | 335 | 0.81 | 579 | 1.41 |
| Tanzania | 73098 | 24.94 | 127035 | 43.35 | 31778 | 10.84 | 52712 | 17.99 |
| Thailand | 10946 | 6.59 | 16341 | 9.84 | 7762 | 4.67 | 13266 | 7.99 |
| The Gambia | 1422 | 45.51 | 1792 | 57.36 | 0 | 0.00 | 0 | 0.00 |
| Timor-Leste | 342 | 8.68 | 757 | 19.21 | 309 | 7.85 | 911 | 23.10 |
| Togo | 808 | 24.85 | 1675 | 51.54 | 413 | 12.72 | 822 | 25.29 |
| Tonga | 0 | 0.00 | 0 | 0.00 | 0 | 0.00 | 0 | 0.00 |
| Trinidad and Tobago | 7 | 0.19 | 12 | 0.32 | 18 | 0.47 | 118 | 3.10 |
| Tunisia | 380 | 4.86 | 468 | 5.99 | 232 | 2.97 | 586 | 7.49 |
| Turkey | 9429 | 7.31 | 14969 | 11.60 | 9394 | 7.28 | 17003 | 13.18 |
| Turkmenistan | 746 | 2.53 | 1071 | 3.63 | 429 | 1.45 | 739 | 2.51 |
| Turks and Caicos Islands | 3 | 1.90 | 5 | 2.87 | 0 | 0.00 | 0 | 0.00 |
| Tuvalu | 0 | 0.00 | 0 | 0.00 | 0 | 0.00 | 0 | 0.00 |
| Uganda | 7605 | 28.83 | 14390 | 54.56 | 3966 | 15.03 | 7864 | 29.81 |
| Ukraine | 97 | 0.34 | 84 | 0.30 | 2992 | 10.53 | 4468 | 15.72 |
| United Arab Emirates | 547 | 2.56 | 780 | 3.65 | 510 | 2.39 | 976 | 4.57 |
| United Kingdom | 1169 | 5.06 | 3358 | 14.55 | 556 | 2.41 | 1373 | 5.95 |
| United States | 67509 | 2.82 | 115931 | 4.84 | 33474 | 1.40 | 58494 | 2.44 |
| United States Minor Outlying Islands | 0 | 0.00 | 0 | 0.00 | 0 | 0.00 | 0 | 0.00 |
| United States Virgin Islands | 0 | 0.00 | 0 | 0.00 | 0 | 0.00 | 0 | 0.00 |
| Uruguay | 662 | 7.96 | 981 | 11.79 | 1305 | 15.69 | 1590 | 19.12 |
| US Naval Base Guantanamo Bay | 0 | 0.00 | 0 | 0.00 | 0 | 0.00 | 0 | 0.00 |
| Uzbekistan | 696 | 1.74 | 701 | 1.76 | 643 | 1.61 | 1201 | 3.01 |
| Vanuatu | 773 | 22.20 | 779 | 22.38 | 270 | 7.76 | 342 | 9.82 |
| Venezuela | 3346 | 0.62 | 3926 | 0.73 | 7014 | 1.30 | 11472 | 2.12 |
| Vietnam | 1493 | 1.14 | 1649 | 1.26 | 4004 | 3.05 | 7301 | 5.56 |
| Wallis and Futuna Islands | 0 | 0.00 | 0 | 0.00 | 0 | 0.00 | 0 | 0.00 |
| Western Sahara | 21 | 0.10 | 59 | 0.27 | 0 | 0.00 | 0 | 0.00 |
| Yemen | 4130 | 5.28 | 6310 | 8.07 | 1877 | 2.40 | 3224 | 4.12 |
| Zambia | 61540 | 22.08 | 110009 | 39.47 | 22247 | 7.98 | 40580 | 14.56 |
| Zimbabwe | 17612 | 22.56 | 29780 | 38.16 | 7463 | 9.56 | 12822 | 16.43 |

| ***Extended Data Table 5.*** *The area and proportion of land identified for conservation and its risk of habitat conversion to human land-use by 2030 and 2050 (following Shared Socioeconomic Pathway 1 (optimistic), 2 (middle of the road), 3 (pessimistic) and an "ensemble" which is the mean of all three) aggregated for developed, emerging and developing countries.* | | | | | | | | | |
| --- | --- | --- | --- | --- | --- | --- | --- | --- | --- |
| **Economic status** | **Total area (km2)** | **Conservation land (km2)** | **Intact Conservation land (km2)** | **Intact Conservation land (%)** | **Shared Socioeconomic pathway** | **Habitat loss by 2030 (km2)** | **Habitat loss by 2030 (%)** | **Habitat loss by 2050 (km2)** | **Habitat loss by 2050 (%)** |
| Developed | 36874471 | 20426643 | 13571807 | 66.44 | SSP1 | 12304 | 0.09 | 49433 | 0.36 |
| Emerging | 58069632 | 27794304 | 20317552 | 73.10 | SSP1 | 48780 | 0.24 | 116659 | 0.57 |
| Developing | 51896702 | 15885454 | 11005400 | 69.28 | SSP1 | 75299 | 0.68 | 154465 | 1.40 |
| Developed |  |  |  |  | SSP2 | 126066 | 0.93 | 221489 | 1.63 |
| Emerging |  |  |  |  | SSP2 | 429945 | 2.12 | 726163 | 3.57 |
| Developing |  |  |  |  | SSP2 | 285427 | 2.59 | 539310 | 4.90 |
| Developed |  |  |  |  | SSP3 | 149065 | 1.10 | 233530 | 1.72 |
| Emerging |  |  |  |  | SSP3 | 345296 | 1.70 | 558695 | 2.75 |
| Developing |  |  |  |  | SSP3 | 786175 | 7.14 | 1393505 | 12.66 |
| Developed |  |  |  |  | Ensemble | 94401 | 0.70 | 165820 | 1.22 |
| Emerging |  |  |  |  | Ensemble | 274671 | 1.35 | 467170 | 2.30 |
| Developing |  |  |  |  | Ensemble | 371523 | 3.38 | 672288 | 6.11 |

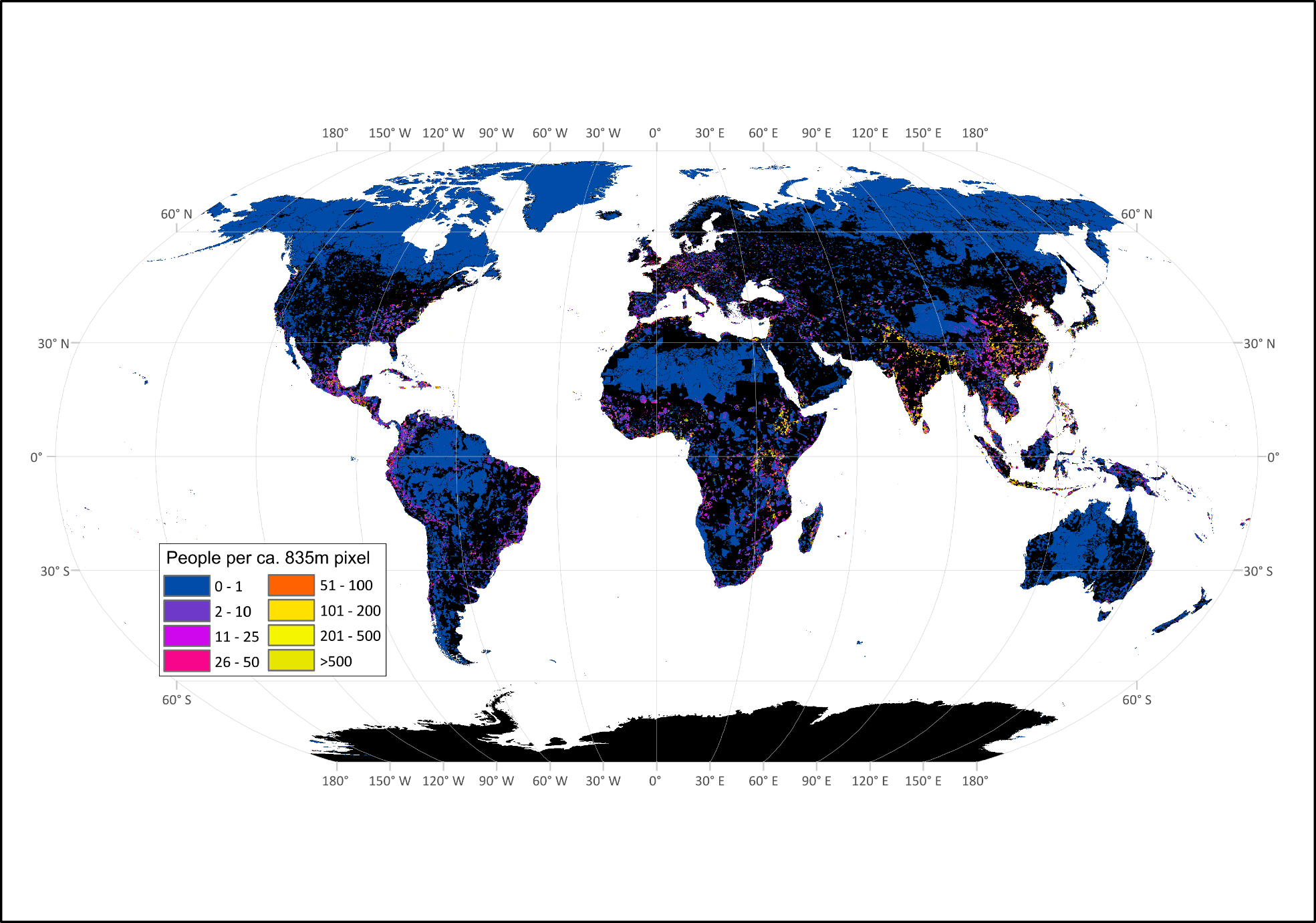

**Extended Data Figure 1.** Human populations living in areas requiring conservation as mapped using LandScan 2018 data at an 835m^2^ resolution.

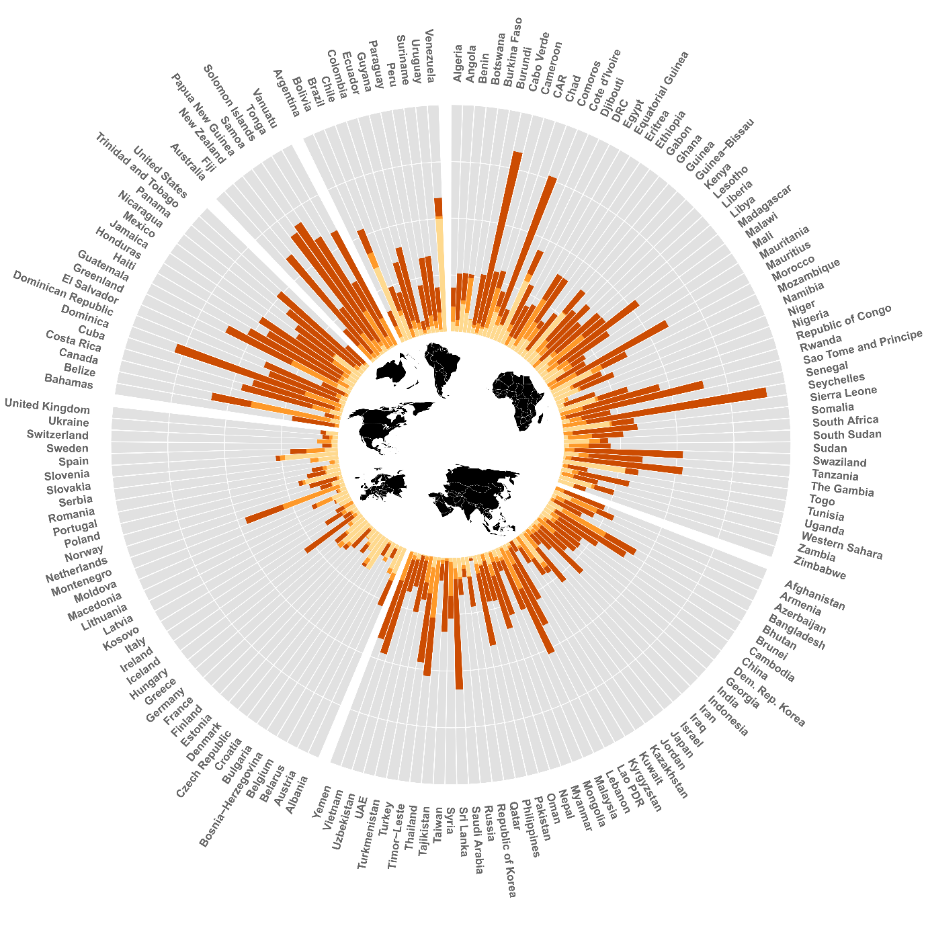

**Extended Data Figure 2.** The proportion of a country’s population living in land requiring conservation. Moving from the centre, the bars represent the percentage of a country’s population in existing protected areas (light orange), Key Biodiversity Areas and ecologically intact areas (medium orange), the additional priority areas (dark orange), and populations not living in important biodiversity conservation areas (grey). The white rings divide the clock graph into four equal parts, each representing 25 percent of a country’s population.

| ***Extended Data Table 6.*** *The area and proportion of intact land requiring conservation within protected areas, Key Biodiversity Areas, and ecologically intact areas (EIA) at risk of habitat conversion by 2030 and 2050 based on Shared Socioeconomic Pathways 1 (optimistic) and 2 (middle of the road), 3 (pessimistic) and "ensemble" which is an average of all three.* | | | | | | | | |
| --- | --- | --- | --- | --- | --- | --- | --- | --- |
| **Economic status** | **Total area (km2)** | **Intact land (km2)** | **Intact land (%)** | **Shared Socioeconomic pathway** | **Habitat loss by 2030 (km2)** | **Habitat loss by 2030 (%)** | **Habitat loss by 2050 (km2)** | **Habitat loss by 2050 (%)** |
| Protected areas | 20482516 | 13960925 | 68.16 | SSP1 | 61117 | 0.44 | 141147 | 1.01 |
| Key Biodiversity Areas | 11618477 | 7304830 | 62.87 | SSP1 | 41328 | 0.57 | 97417 | 1.33 |
| EIA | 35151586 | 29076607 | 82.72 | SSP1 | 10254 | 0.04 | 52174 | 0.18 |
| Protected areas |  |  |  | SSP2 | 364826 | 2.61 | 638106 | 4.57 |
| Key Biodiversity Areas |  |  |  | SSP2 | 256808 | 3.52 | 450589 | 6.17 |
| EIA |  |  |  | SSP2 | 173397 | 0.60 | 319755 | 1.10 |
| Protected areas |  |  |  | SSP3 | 577952 | 4.14 | 998687 | 7.15 |
| Key Biodiversity Areas |  |  |  | SSP3 | 402411 | 5.51 | 676934 | 9.27 |
| EIA |  |  |  | SSP3 | 206129 | 1.48 | 358069 | 2.56 |
| Protected areas |  |  |  | Ensemble | 334632 | 2.40 | 592647 | 4.25 |
| Key Biodiversity Areas |  |  |  | Ensemble | 233515 | 3.20 | 408313 | 5.59 |
| EIA |  |  |  | Ensemble | 129927 | 0.45 | 243333 | 0.84 |

| ***Extended Data Table 7.*** *Table shows the area (millions of km^2^) and (percent) of overlap between priority areas identified using different cost metrics. Antarctica is excluded from area calculations since it is beyond the extent of the human footprint and agricultural opportunity data.* | | | | | | |
| --- | --- | --- | --- | --- | --- | --- |
|  | Area | Agricultural opportunity | Sum human footprint | Area | Agricultural opportunity | Sum human footprint |
| *Excluding urban* |  |  |  |  |  |  |
| Area |  | 8.8 (74.2) | 8.6 (72.6) | 6.5 (55.2) | 5.7 (48.3) | 5.6 (47.4) |
| Agricultural opportunity | 8.8 (52.5) |  | 9.8 (58.6) | 6.2 (37.1) | 11.8 (70.8) | 6.7 (40.2) |
| Sum human footprint | 8.6 (65.8) | 9.8 (75.2) |  | 6.1 (46.8) | 6.9 (52.9) | 8.6 (65.8) |
| *Excluding agriculture and urban* |  |  |  |  |  |  |
| Area | 6.5 (58.6) | 6.2 (55.7) | 6.1 (54.9) |  | 7.7 (69.5) | 7.9 (71.4) |
| Agricultural opportunity | 5.7 (36.3) | 11.8 (75.2) | 6.9 (43.8) | 7.7 (49.2) |  | 8.6 (54.8) |
| Sum human footprint | 5.6 (46.2) | 6.7 (55.5) | 8.6 (70.9) | 7.9 (65.6) | 8.6 (71.2) |  |

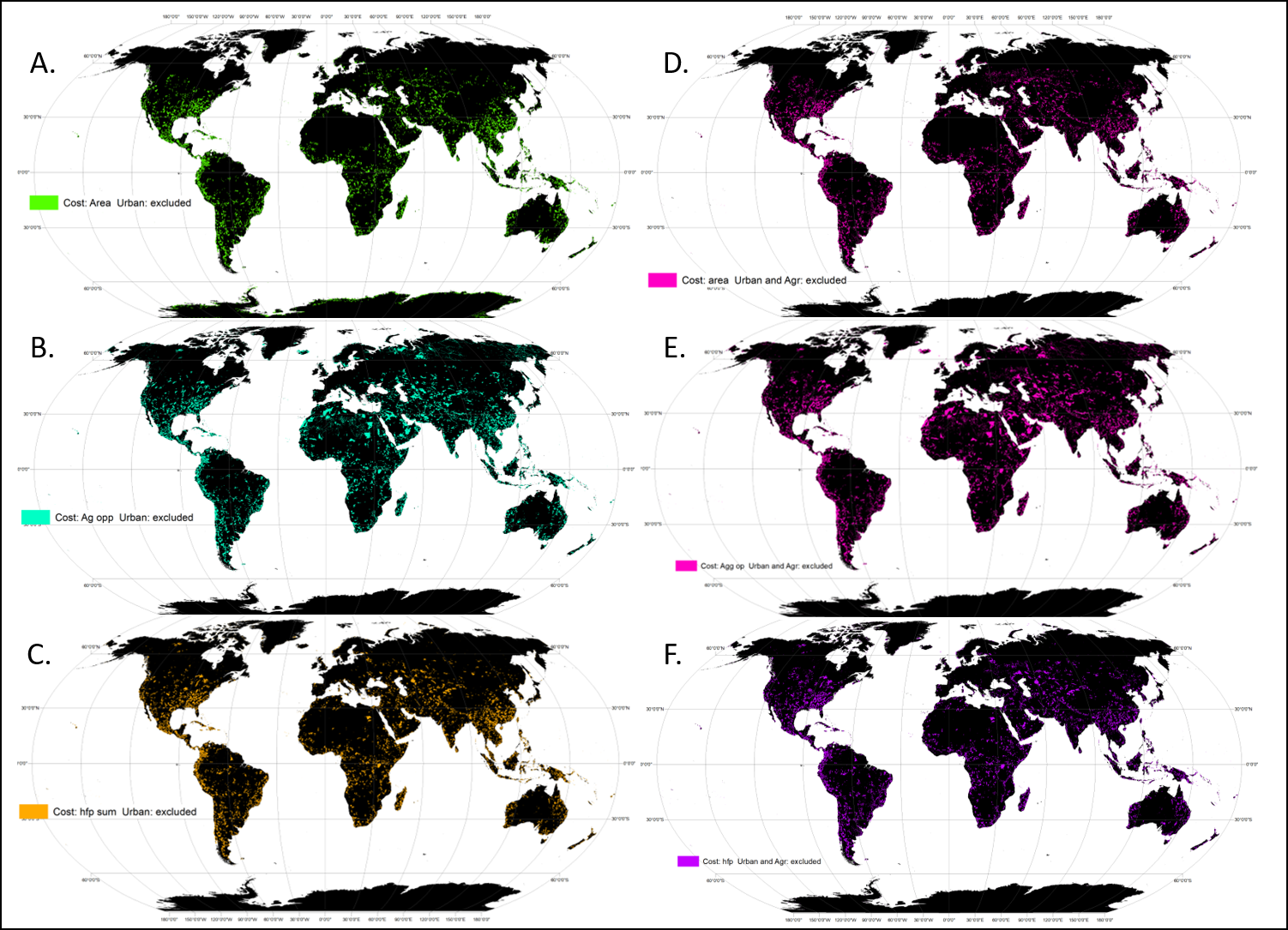

**Extended Data Figure 3.** Additional priority areas under different cost and current land-use scenarios. Including A) where the cost is planning unit area and urban areas are excluded, B) where the cost is agricultural opportunity and urban areas are excluded, C) where the cost is the sum of the human footprint and urban areas are excluded, D) where the cost is area and both urban and agricultural areas are excluded, E) where the cost is agricultural opportunity and both urban and agricultural areas are excluded, and F) where the cost is the sum of the human footprint and both urban and agricultural areas are excluded. Note that in D,E,F when agricultural areas are excluded, it is not possible to meet all species targets.

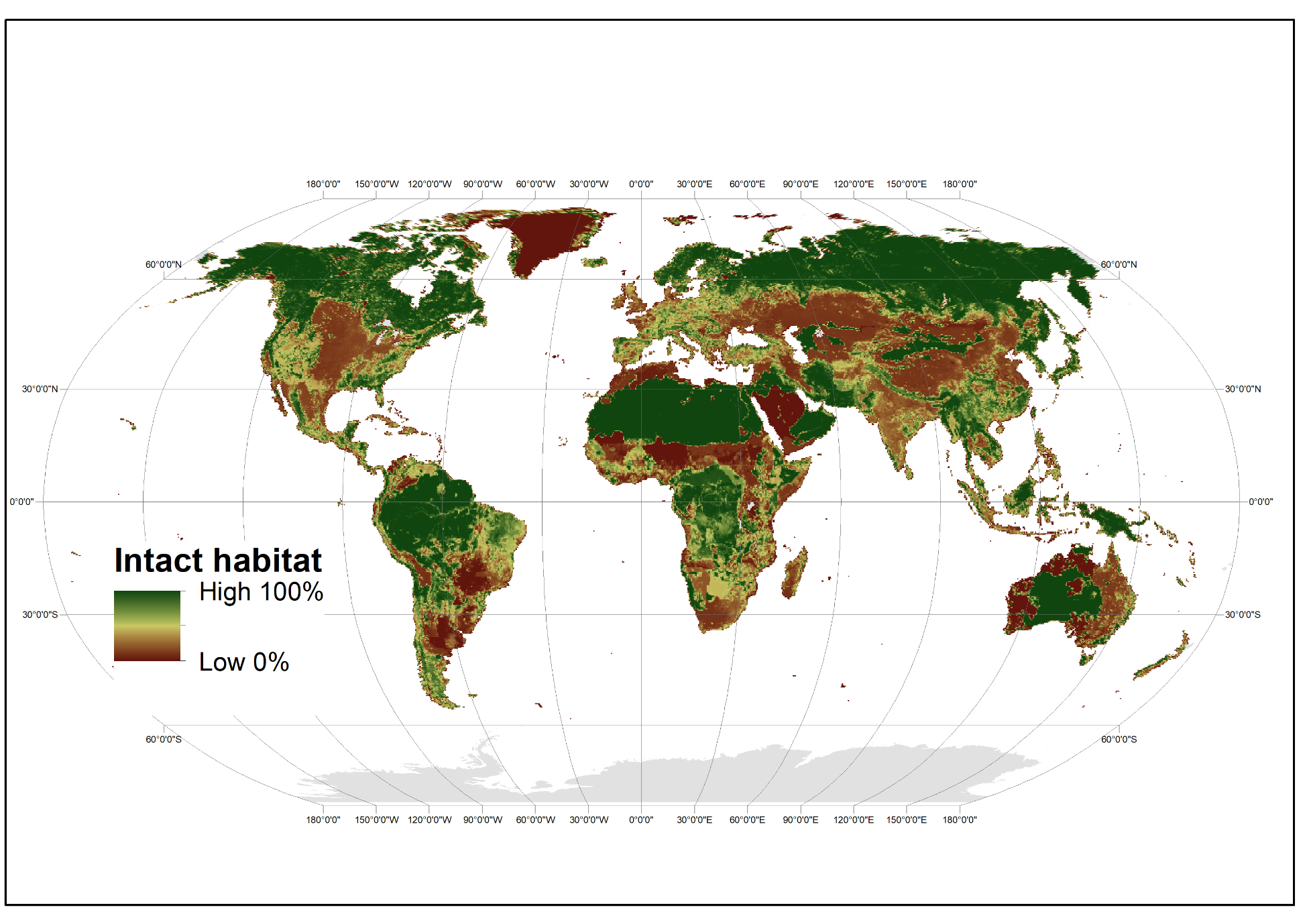

**Extended Data Figure 4.** Proportion of intact natural habitat in each planning unit.

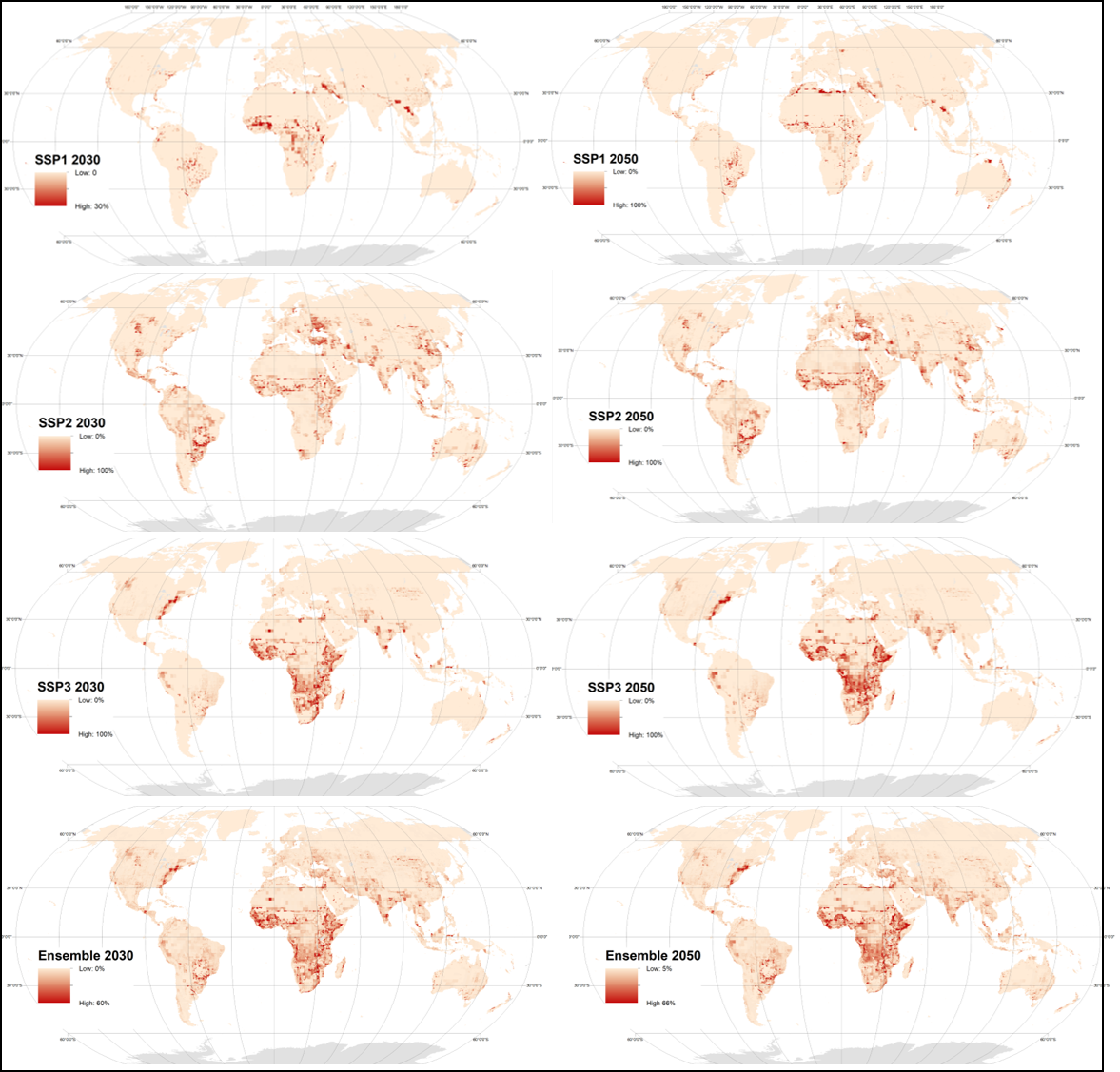

**Extended Data Figure 5.** Projected natural habitat conversion under Shared Socioeconomic Pathways 1, 2, 3, and an ensemble (average of them all) for the years 2030 and 2050.
